# Neuronal lineage tracing from progenitors in human cortical organoids reveals novel mechanisms of human neuronal production, diversity, and disease

**DOI:** 10.1101/2023.06.17.545314

**Authors:** Luke A.D. Bury, Shuai Fu, Anthony Wynshaw-Boris

## Abstract

The contribution of progenitor subtypes to generate the billions of neurons during human cortical neurogenesis is not well understood. We developed the Cortical ORganoid Lineage Tracing (COR-LT) system for human cortical organoids. Differential fluorescent reporter activation in distinct progenitor cells leads to permanent reporter expression, enabling the progenitor cell lineage of neurons to be determined. Surprisingly, nearly all neurons produced in cortical organoids were generated indirectly from intermediate progenitor cells. Additionally, neurons of different progenitor lineages were transcriptionally distinct. Isogenic lines made from an autistic individual with and without a likely pathogenic variant in the CTNNB1 gene demonstrated that the variant substantially altered the proportion of neurons derived from specific progenitor cell lineages, as well as the lineage-specific transcriptional profiles of these neurons, suggesting a pathogenic mechanism for this mutation. These results suggest individual progenitor subtypes play unique roles in generating the diverse neurons of the human cerebral cortex.

**Graphic Abstract.**
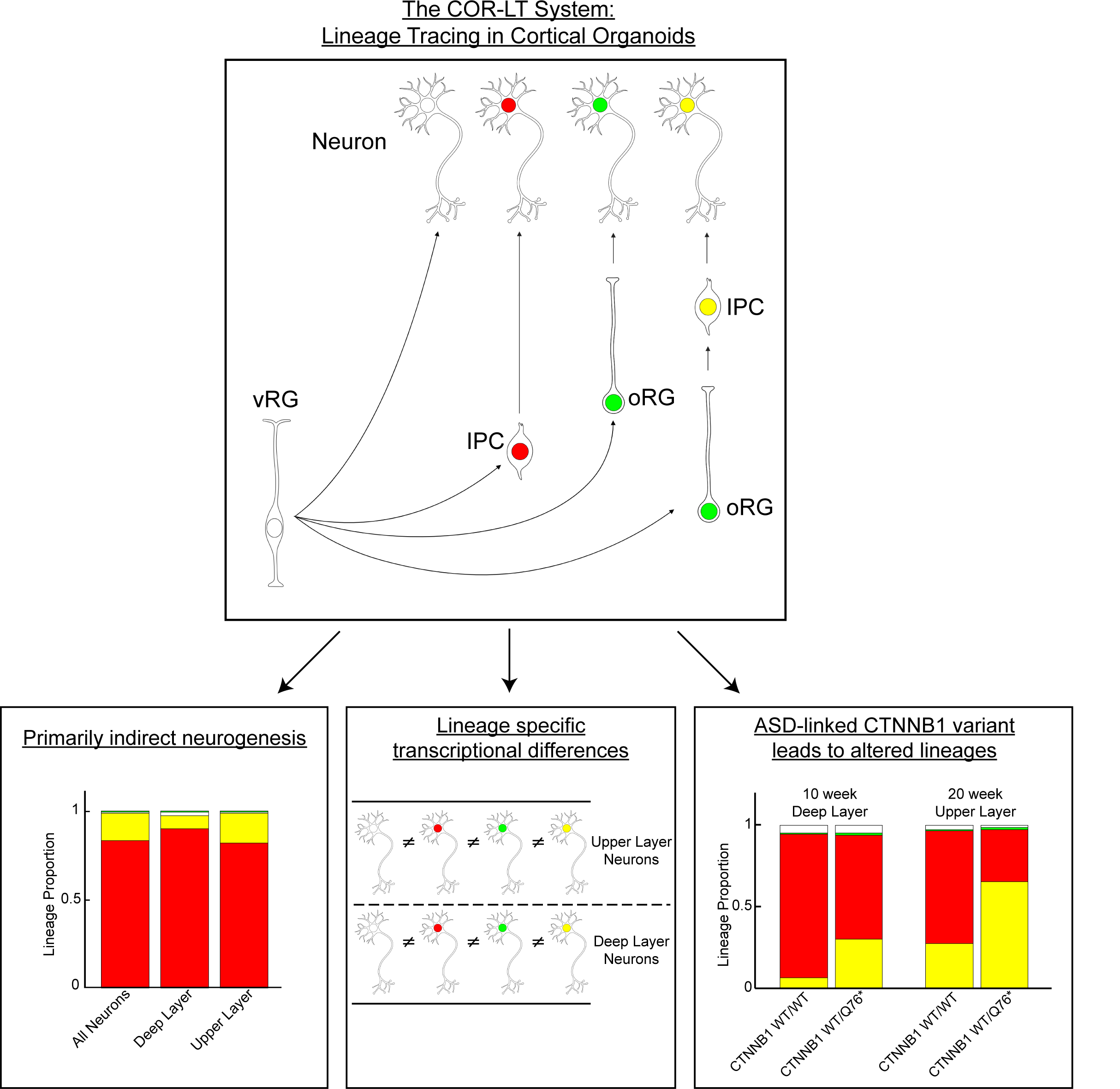

## Highlights

1. The Cortical ORganoid Lineage Tracing (COR-LT) system records the genetically-defined progenitor cell lineage of neurons in human cortical organoids.
2. Almost all neurons in human cortical organoids are produced indirectly through intermediate progenitor cells.
3. Neurons from different lineages are transcriptionally distinct.
4. An autism spectrum disorder-specific *CTNNB1* stop-gain variant leads to altered lineage-specific neuronal production and transcriptional differences.

## Introduction

During mid-fetal development, over the course of five to six months, nearly all of the approximately 16 billion neurons of the cerebral cortex are generated. Producing the correct number and types of cortical neurons during this time period is essential to produce a properly functioning brain, as illustrated by the severe phenotypes accompanying the extensive neuron loss associated with microcephaly,^1, 2^ and more subtle alterations in cortical neuron number found in a variety of neurodevelopmental disorders, including autism spectrum disorder (ASD).

In the human cortex, glutamatergic neurons are the main excitatory neuronal population and make up a majority of the total number of cortical neurons. These neurons are produced in the dorsal telencephalon during mid-gestation through a variety of neural progenitor cell (NPC) subtypes including ventricular radial glial (vRG) cells,^3, 4^ outer radial glial (oRG) cells^5–8^ and transient-amplifying intermediate progenitor cells (IPCs).^9–16^ These transcriptionally and morphologically unique NPC subtypes can potentially self-replicate, produce other progenitor subtypes, or undergo neurogenic divisions to produce the diverse types of glutamatergic neurons found in the cerebral cortex.

The lineage of cortical neurons from these NPC subtypes plays an important role in mammalian brain development, demonstrated by genetic lineage tracing systems in mice.^14–21^ However, the extent to which each progenitor subtype contributes to the final neuronal population in cortical organoids and in the developing human brain is unknown. It is also not known whether neurons produced from different progenitors possess any meaningful biological differences in humans. Finally, the extent to which neurodevelopmental diseases perturb neurogenesis in specific progenitor subtypes and/or alter any unique biological characteristics that might be found in specific NPC-subtype-derived neurons is unclear. For example, subtle alterations in neurogenesis may be difficult to detect in ASDs, where the brain is structurally normal but may display neuron number differences when associated with macrocephaly or microcephaly.

In order to determine the proportion and detailed characterization of neuron populations produced by specific NPC subtypes, we developed a system to record neuronal lineage in human cortical organoids (COR-LT – Cortical ORganoid Lineage Tracing). In the COR-LT system, the NPC-subtype-enriched genes *EOMES* (*TBR2)* in IPCs^22^ and *HOPX* in oRGs^23, 24^ drive the expression of unique DNA recombinases. This subsequently leads to fluorescent protein expression in the progenitors themselves and the neuronal progeny that they produce. We used this system to directly determine the proportion of neurons generated by individual NPC subtypes in cortical organoids. Combining the COR-LT system with single-cell RNA-sequencing (scRNA-seq) led to the detection of transcriptomic differences in similar neuronal populations that were generated via different progenitor subtypes. Finally, we employed this system in an isogenic model of ASD to track how both neuronal production from different NPC subtypes and the transcriptomic characteristics of neurons derived from unique NPC subtypes were altered by a single genetic variant.

## Results

### The COR-LT system design

To implement and validate the COR-LT system, four unique DNA cassettes were sequentially incorporated into the genome of a neurotypical control induced pluripotent stem (iPS) cell line (“Clay” from ^25^) via CRISPR/Cas9 and TALEN genome editing (Fig. 1A, B). A DNA sequence encoding T2A-linked Cre recombinase was incorporated into the 3’ end of the endogenous *EOMES* gene (Fig. 1A), while a T2A-linked HA-tagged Dre recombinase was incorporated into the 3’ end of the endogenous *HOPX* gene (Fig. 1B). Additionally, two unique “stop-flox/rox” reporter constructs were incorporated into separate “safe harbor” genomic sites. In these reporter constructs, the promoter and reporter are separated by a loxP- or roxP-flanked STOP sequence which prevents reporter expression unless the STOP sequence is cleaved by either Cre or Dre recombinase, respectively. Specifically, a CAG-promoter-driven “stop-flox” tdTomato reporter with a nuclear localization sequence (NLS) was integrated into the AAVS1 safe harbor site^26, 27^ (Fig. 1A), while a similar “stop-rox” FLAG-tagged ZsGreen-NLS reporter construct was integrated into the H11 safe harbor site^28^ (Fig. 1B).

**Figure 1.**
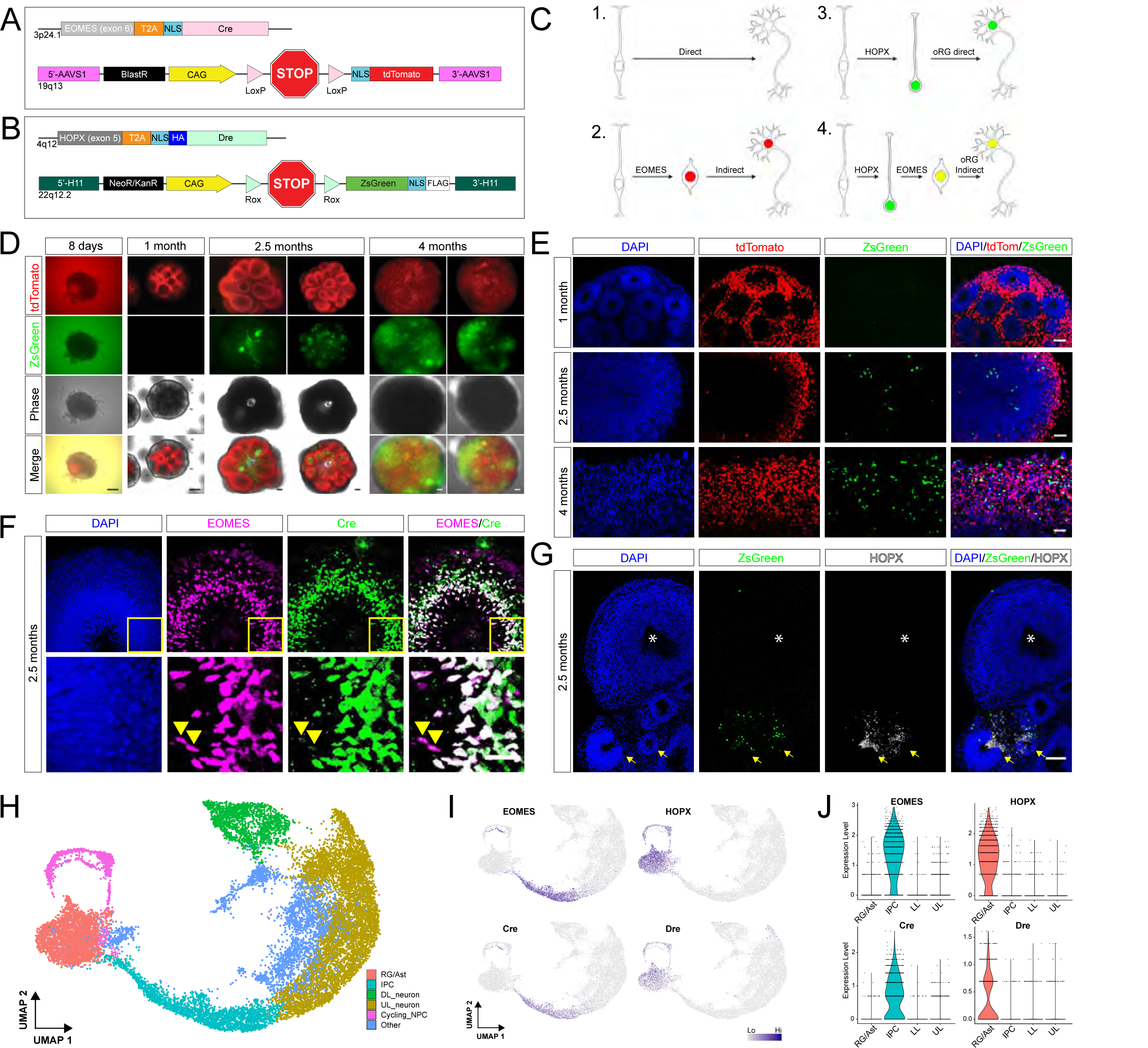
COR-LT system design and verification. (A, B) Constructs used for genome editing. (C) Diagram of possible lineages that can be identified with the COR-LT system. (D, E) Time-course of tdTomato (red) and ZsGreen (green) expression in control COR-LTs organoids in live (D) and fixed (E) tissue. (F) Immunohistochemical (IHC) detection of EOMES (magenta) and Cre (green) in a 2.5 month-old control COR-LT organoid section. Yellow box in top row indicates region shown in bottom row. Arrowheads indicate EOMES+ cells with faint Cre immunoreactivity. (G) IHC detection of HOPX (white) with ZsGreen expression (green) in a 2.5 month-old organoid section. Arrows indicate HOPX+/ZsGreen+ progenitor regions. Astrisk indiates a HOPX-/ZsGreen-progenitor region. (H) UMAP of integrated control scRNA-seq experiments with general cell classifications. (I) UMAPs showing expression of COR-LT components in individual cells. (J) Violin plots showing expression of COR-LT components within specific cell groups. Scale bars: (D) = 200um, (E), (F) = 25um, (G) = 100um. (C) Created with BioRender.com.

In this system, expression of *EOMES* in IPCs or *HOPX* in oRGs leads to the co-expression of either Cre or Dre recombinase, respectively. Recombinase expression then leads to the removal of stop sequences in the tdTomato or ZsGreen reporters, enabling permanent expression of the reporter genes in the nuclei of cells that expressed *EOMES* or *HOPX*, respectively. Due to the CAG constitutive promoter, these fluorescent reporters will continue to be expressed in all daughter cells, including both progenitors and neurons, that are derived from the cell with the original recombination event (Fig. 1C).

The COR-LT system was designed to identify the lineage of four types of neurons within the organoid: (1) neurons that do not express a fluorescent reporter arise from direct vRG neurogenesis (vRG->neuron); (2) neurons that express tdTomato only arise from an IPC via indirect neurogenesis (vRG->IPC->neuron); (3) neurons that express ZsGreen only are generated via direct oRG neurogenesis (oRG->neuron); and (4) neurons expressing both tdTomato and ZsGreen would have a lineage that progressed through both an oRG and IPC prior to neuronal differentiation via indirect neurogenesis (oRG->IPC->neuron) (Fig. 1C). We next verified the lineage of each of these types of neurons based on the expression of these fluorescent reporters and the recombinases that activate the system.

### Verification of appropriate COR-LT system activation

To properly characterize neuronal lineage in the organoid, it is critical that the recombinases and reporters are only expressed in the appropriate progenitor cells and neurons. In control iPS cells, neither *EOMES* nor *HOPX* were expressed at significant levels, resulting in minimal reporter expression (<0.5% of cells, Supplemental Fig. 1A, data not shown). To prevent even minimal inappropriate reporter expression in organoids, iPS colonies expressing either reporter were identified via fluorescent microscopy and removed from culture prior to each organoid experiment. While endogenous reporter activation was low in iPS cells, transient transfection of DNA plasmids encoding either Cre or Dre recombinase led to specific tdTomato or ZsGreen expression, respectively (Supplemental Fig. 1A).

Cortical organoids were produced from control iPS cells with the COR-LT system following a modified version of previously established protocols^29–31^ and the time course of reporter activation in live (Fig. 1D) and fixed (Fig. 1E) samples was established. During early organoid development, neither *EOMES*+ IPCs nor *HOPX*+ oRG cells were produced, resulting in a lack of reporter expression in the organoid (Fig. 1D). By 1 month, expression of the tdTomato reporter was found surrounding the numerous ventricular zone-like structures in these organoids (Fig. 1D, E), coincident with the location of *EOMES*+ IPCs (Fig. 1F). There was a lack of ZsGreen+ cells within the organoids at 1 month (Fig. 1D, E), before *HOPX*+ oRG cells were present. After 2.5 months of development, the organoids had grown larger and the number of tdTomato+ cells had increased, while the initial generation of ZsGreen+ cells was observed (Fig. 1D, E), corresponding to HOPX+ oRG cell production (Fig. 1G). Finally, at four months, a substantial increase in the number of tdTomato+ and ZsGreen+ cells was observed, consistent with an expansion of the numbers of both IPCs and oRGs, along with their neuronal progeny (Fig. 1D, E).

To ensure that reporter expression does not adversely affect organoid development, iPS cells from the control COR-LT system cell line were transiently co-transfected with Cre and Dre recombinases. Individual iPS cell clones were isolated, and clones with 100% co-activation of both reporters (tdTomato and ZsGreen) were used to generate organoids. Organoids grown to 9-weeks produced well-defined ventricular-like regions along with neurons outside of these progenitor regions (Supplemental Fig. 1B), indicating that long-term co-expression of both reporters in individual cells does not exert major effects on organoid health. Additionally, both reporters were detected in a large majority of TBR1+ neurons, indicating minimal reporter silencing (98.8% tdTomato+; 93.2% ZsGreen+, Supplemental Fig. 1B, C).

To further validate recombinase expression in appropriate cell types, 10-week-old organoid sections were co-labeled with antibodies detecting Cre recombinase and EOMES proteins, which indicated nearly all EOMES+ cells appropriately expressed Cre recombinase (Fig. 1F). At 10 weeks of organoid development, the production of oRG cells commenced with some progenitor regions expressing HOPX and some progenitor regions lacking HOPX expression. Immunohistochemistry (IHC) of 10-week organoid sections revealed that only progenitor regions that contained HOPX+ cells were also positive for ZsGreen, suggesting appropriate activation of the ZsGreen reporter in oRG cells within the organoid (Fig. 1G). Co-labeling with antibodies detecting Dre recombinase were unsuccessful, as was detection of the HA sequence that was added to the 5’ end of the Dre construct, possibly due to the 3’ residues of the T2A sequence obscuring the HA epitope.

To further validate the COR-LT system in these control organoids, single cell RNA-sequencing (scRNA-seq) was performed on four-month-old organoids consisting of two biological replicates from independent experiments, which were integrated together via Seurat (Fig. 1H, Supplemental Fig. 2A). Similar cell types were generated in each experiment, including astroglial progenitor cells (“RG/Ast”), intermediate progenitor cells (“IPC”), cycling progenitor cells (“Cycling_NPC”) and neurons from both upper (“UL_neuron”) and deep (“DL_neuron”) cortical layers (Fig. 1H, Supplemental Fig. 3A, B, Supplemental Table 1, https://github.com/lbury3/cor_lt/). Cells identified as “Other” were enriched for mitochondrial genes, lncRNAs, ribosomal genes, and/or the lncRNA gene *MALAT1* (Supplemental Table 1, https://github.com/lbury3/cor_lt/) and excluded from downstream analysis.

The recombinases of the COR-LT system were also detected via scRNA-seq. As expected, Cre recombinase mRNA was found almost exclusively in IPCs (Fig. 1I, J) and was only significantly enriched in clusters where *EOMES* was also significantly enriched (Supplemental Table 1, https://github.com/lbury3/cor_lt/). Likewise, Dre recombinase was detected almost exclusively in astroglial progenitor cells (Fig. 1I, J) and only in clusters where *HOPX* was significantly enriched (Supplemental Table 1, https://github.com/lbury3/cor_lt/). Thus, both IHC and scRNA-seq data validate the activation of the COR-LT system exclusively in cells derived from IPC and oRG lineages.

### Nearly all neurons in control cortical organoids were produced indirectly through IPCs

In mice, up to 60-70% of neurons are produced indirectly from IPCs.^14, 19^ However, the proportion of neurons produced indirectly through IPCs in the human brain or in human model systems is unclear. Therefore, scRNA-seq analysis of four-month-old COR-LT system control organoids was employed to determine the proportion of IPC-derived tdTomato+ neurons in both deep layer and upper layer neuronal clusters.

Remarkably, tdTomato was expressed in practically all neurons, including both upper layer (UL) and deep layer (DL) groups (Fig. 1H, Fig. 2A, C, Supplemental Fig. 5A; DL = 98.0% (1727/1762); UL = 99.5% (6412/6442)). This transcriptomic data was supported by IHC in organoid tissue sections, which indicated nearly 100% co-localization of tdTomato with the upper layer neuronal marker SATB2 and the deep layer neuronal marker TBR1 (Fig. 2D-F, SATB2+/tdTomato+ = 98.4% +/− 2.3; TBR1+/tdTomato+ = 98.6% +/− 1.4, n = 10 organoid sections), demonstrating that almost all neurons produced in these four-month-old control organoids were indirectly generated through intermediate progenitor cells.

**Figure 2.**
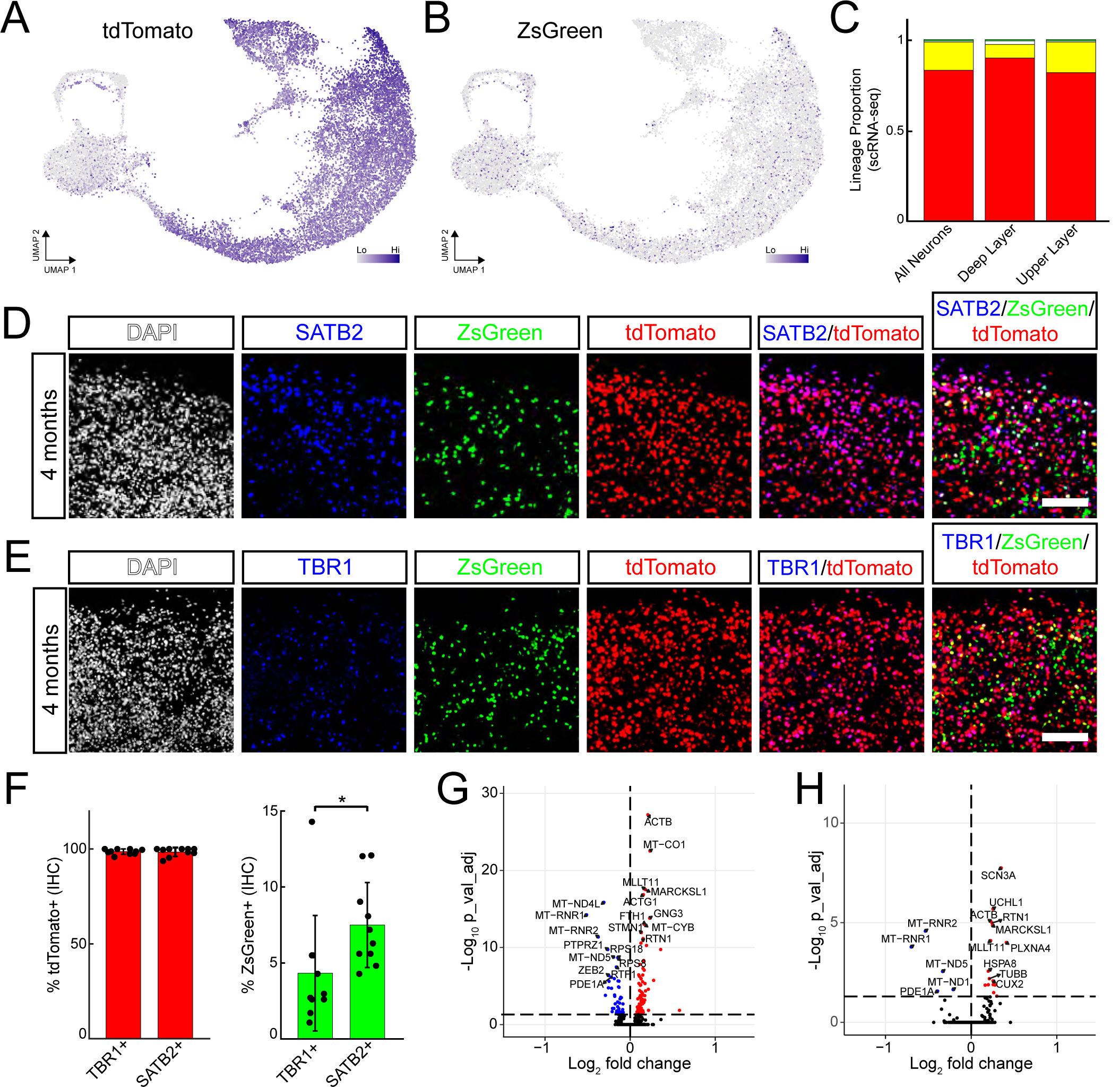
The COR-LT system indicates primarily indirect neurogenesis in cortical organoids and differential gene expression in upper layer neurons of distinct lineages. (A, B) UMAPs of tdTomato and ZsGreen expression as determined by scRNA-seq in 4-month-old control COR-LT organoids. (C) Lineage proportions as determined by reporter expression in scRNA-seq data in 4-month-old control COR-LT organoids. (D, E) IHC detection of SATB2+ upper layer neurons (D, blue) or TBR1+ deep layer neurons (E, blue) with ZsGreen (green) and tdTomato (red) expression in 4-month-old control COR-LT organoid sections. Scale bars = 100um. (F) IHC quantification (n = 10 organoid images per group, error bars = SD, *p<0.05) (G) Volcano plot demonstrating genes significantly enriched (p_val_adj < 0.05) in “yellow” lineage upper layer neurons (right side, red) or “red” lineage upper layer neurons (left side, blue) in four-month-old control COR-LT organoids. (H) Volcano plot demonstrating genes significantly enriched (p_val_adj < 0.05) in “yellow” lineage upper layer neurons from cluster 1 (right side, red) or “red” lineage upper layer neurons from cluster 1 (left side, blue) in four-month-old control COR-LT organoids. For volcano plots, horizontal dashed line = p_val_adj = 0.05; vertical dashed line = 0.

It is possible that sparse Cre recombinase expression in astroglial cells or neurons themselves could lead to an accumulation of tdTomato+ neurons without going through the IPC lineage. However, both *EOMES* and Cre recombinase were almost exclusively expressed together in IPCs, as indicated by scRNA-seq (Fig. 1I, J). In neurons especially, where individual neurons can exist within the organoid for long periods of time, it is possible that even sporadic, transient Cre recombinase expression could lead to an accumulation of tdTomato+ cells over longer periods of culture. However, in 5-week-old organoids, which contain an inherently large number of newborn neurons, practically all neurons were tdTomato+ (Supplemental Fig. 4A). Together, these findings strongly suggest that initial tdTomato activation in the COR-LT system was specific to IPCs expressing *EOMES*-driven Cre recombinase and that these cells were the primary source of neurons in control organoids.

### Neurons derived from oRGs vs vRGs in control organoids were transcriptionally distinct

Lineage proportion analysis of scRNA-seq data indicated that 15.3% (1257/8204) of all neurons expressed ZsGreen at four months in control organoids (Fig. 1H, Fig. 2B, C, Supplemental Fig. 5B). All but three of these cells also express tdTomato (1254/1257, 99.8%), indicating that oRG->IPC->neuron indirect neurogenesis predominated over oRG->neuron direct neurogenesis. A higher percentage of upper layer neurons were produced through oRG progenitors (Fig. 2C, 17.4%, 1120/6442) compared to deep layer neurons (Fig. 2C, 7.8%, 137/1762), reflecting the role of oRGs in later-stage, upper layer neurogenesis.^5^ Significant enrichment of ZsGreen in upper layer neurons compared to deep layer neurons was also confirmed via IHC (Fig. 2D-F, SATB2+/ZsGreen+ = 7.5% +/− 2.8; TBR1+/ZsGreen+ = 4.3% +/− 3.8, n = 10 organoid sections, p = 0.0477). The majority of the remaining neurons were tdTomato+/ZsGreen-indicating they were produced via a vRG->IPC->neuron lineage (Fig. 2C, UL neurons = 82.2%, (5294/6442); DL neurons = 90.3% (1591/1762)).

It is not known whether neurons of similar subtypes (e.g. same cortical layer) produced from an oRG are distinct from neurons that are not oRG-derived in either mouse or human. Since oRG-derived ZsGreen+ neurons were enriched in the upper layer group, we directly compared the transcriptional profiles of vRG->IPC derived upper layer neurons that expressed tdTomato but not ZsGreen (“red” lineage) and oRG->IPC derived upper layer neurons that expressed both tdTomato and ZsGreen (“yellow” lineage). This comparison identified 51 genes that were upregulated in vRG->IPC derived “red” lineage neurons, including eight SFARI autism risk genes (HIVEP2, ANK2, ANP32A, ZBTB18, AFF2, HNRNPR, NCKAP5, and SYT1), while 90 genes were upregulated in oRG->IPC derived “yellow” lineage neurons (minus reporter genes themselves), including seven known SFARI autism risk genes (ACTB, CHST2, EEF1A2, TSPAN7, ARF3, GNB2, and GNAS) (Fig. 2G, Supplemental Table 2, https://github.com/lbury3/cor_lt/). Importantly, there were no significant differences in gene expression between two groups of randomly selected UL neurons (Supplemental Fig. 5D).

To gain further biological insight into the molecular distinctions of oRG- and vRG-derived neurons, Gene Set Enrichment Analysis (GSEA) was performed, indicating upregulation of a single gene ontology (GO) term (“cytoplasmic translation”) in “red” lineage neurons compared to “yellow” lineage neurons (Supplemental Fig. 5C). Additional GSEA analysis demonstrated an upregulation of synapse-related GO terms in “red” and “yellow” lineage upper layer neurons compared with direct vRG-derived non-expressors of either tdTomato or ZsGreen (“no color” lineage) upper layer neurons (Supplemental Fig. 5C, Supplemental Table 3, https://github.com/lbury3/cor_lt/). However, this analysis might be confounded based on the relatively low numbers of neurons that did not express either reporter. The number of neurons expressing ZsGreen but not tdTomato (“green”) were not sufficient to perform similar GSEA comparisons.

Gene expression differences between “red” and “yellow” lineage neurons in individual UL scRNA-seq clusters were also determined (Supplemental Fig. 3A, clusters 1, 3, 4, 7, 8, 15, 18, 19). Neurons in cluster 1 displayed the highest number of differentially expressed genes compared to other UL neuronal clusters, with *SCN3A* being the top enriched gene in oRG-derived “yellow” lineage neurons (Fig. 2H, Supplemental Table 4, https://github.com/lbury3/cor_lt/). Mutations in *SCN3A*, which codes for the voltage-gated sodium channel subunit Na_v_1.3, cause a distinct neurodevelopmental disorder characterized by seizures, cortical malformations, speech deficits, and intellectual disability.^32^ SCN3A was enriched in the outer subventricular zone and cortical plate of human fetal brain,^32^ suggesting that SCN3A is enriched in oRG cells and neurons that are potentially derived from these progenitors. *SCN3A* up-regulation in ZsGreen+ neurons provides direct evidence of this lineage progression in organoids and suggests the COR-LT system can identify populations of neurons in cortical organoids that reflect biologically relevant neuronal populations of the developing human brain.

### CTNNB1 p.Gln76* lead to delayed neuron production in ASD cortical organoids

In addition to assessing aspects of normal development, the COR-LT system has the potential to elucidate novel disease mechanisms related to cortical neuron production, especially when alterations in neurogenesis may be subtle, such as ASDs associated with macrocephaly.^33–37^ Therefore, the system was used to characterize organoids generated from a previously described iPS cell line derived from an individual with ASD and macrocephaly which contains a heterozygous CTNNB1 p.Gln76* variant (“Arch” as described in ^25^). This specific variant likely contributes to ASD pathology as the majority of *CTNNB1* mutations identified in individuals with ASD are stop-gain variants (https://gene.sfari.org/database/human-gene/CTNNB1#variants-tab).

To assess the effect that this variant has on cortical organoid development, the four COR-LT system components were integrated into the patient line with the CTNNB1 p.Gln76* variant. The CTNNB1 p.Gln76* variant was then corrected to WT via genome editing after the four COR-LT system components were integrated into the patient line (see Methods section). Thus, isogenic COR-LT system cell lines were generated with both the patient-derived *CTNNB1* mutation (CTNNB1 WT/Q76*) and a “corrected” version of this mutation (CTNNB1 WT/WT) enabling the developmental effect of the likely-pathogenic *CTNNB1* mutation to be assessed without confounding genetic background differences.

Cortical organoids were produced from both cell lines using a slightly modified protocol compared to control organoids (see Methods). “Leaky” tdTomato in the iPS cell stage was detected at a higher rate in both isogenic ASD cell lines compared to control, but constituted <5% of all cells total. scRNA-seq experiments were conducted at week 10 and week 20 of organoid development. Cortical organoids from both cell lines produced expected NPC populations including astroglial progenitors, IPCs, and cycling progenitor cells as well as neuronal populations that included deep (10-week, 20-week) and upper layer neurons (20-week) in addition to a small number of interneurons (Supplemental Fig. 6A-D, Supplemental Table 5, https://github.com/lbury3/cor_lt/). Datasets from the two cell lines integrated well at both weeks 10 and 20 (Supplemental Fig. 2B, C), reflecting similar cell types from both organoid genotypes.

Proper function of the system was verified in a similar fashion to the COR-LT system in control organoids (Fig. 1). Appearance of the COR-LT system fluorescent reporters occurred on approximately the same time-course as in control, with the notable exception of ZsGreen production in CTNNB1 WT/Q76* organoids at 5 weeks (Fig. 3A, B). IHC co-labeling with EOMES and Cre antibodies indicated that practically all EOMES+ cells also expressed Cre recombinase (Supplemental Fig. 7A), while ZsGreen+ cells were only produced in organoids that expressed HOPX (Fig. 3C). In addition, scRNA-seq data indicated that *EOMES* and Cre recombinase expression was restricted to IPCs while *HOPX* and Dre recombinase expression was restricted to astroglial progenitors (Supplemental Fig. 6A-D, Supplemental Fig. 8A-D).

**Figure 3.**
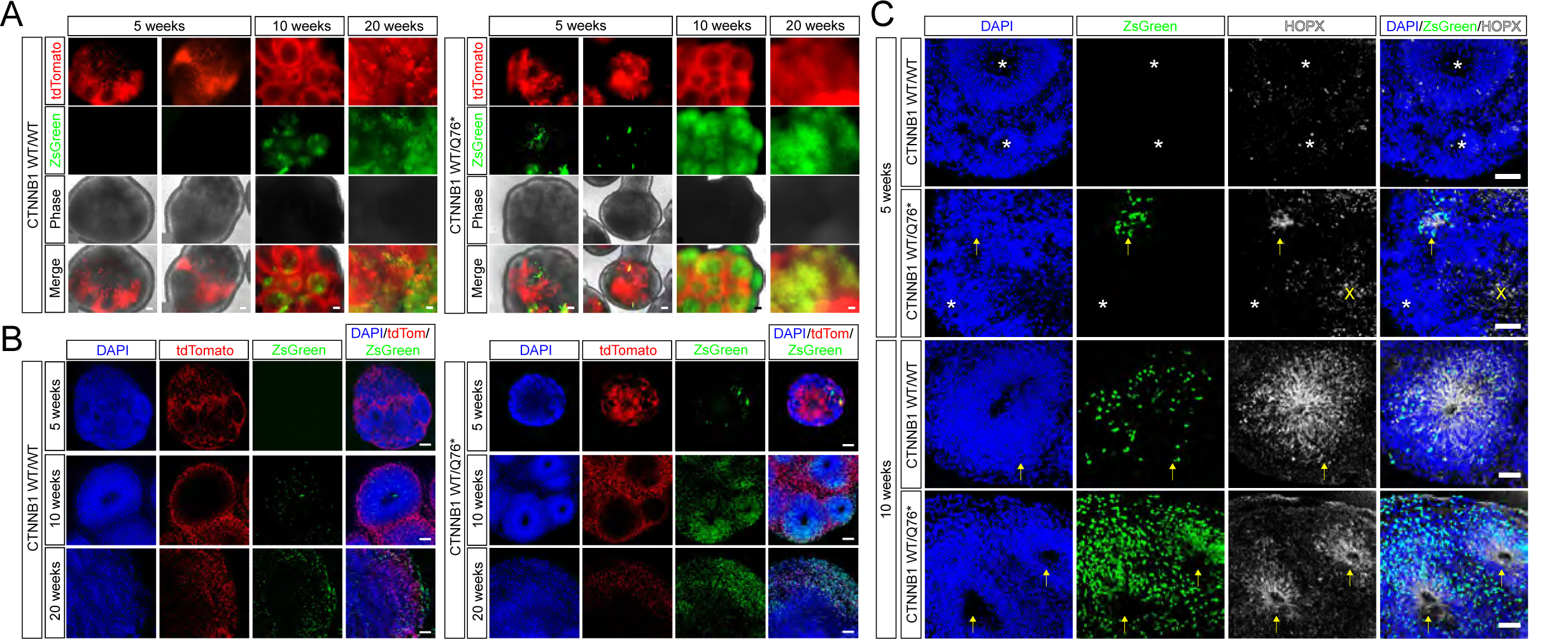
The COR-LT system detects early oRG production in CTNNB1 WT/Q76* organoids. (A, B) Time-course of tdTomato (red) and ZsGreen (green) expression in CTNNB1 WT/WT (left) and CTNNB1 WT/Q76* (right) organoids in live (A) and fixed (B) tissue. (C) IHC detection of HOPX (white) with ZsGreen expression (green) in 5-week-old (top) and 10-week-old (bottom) ASD COR-LT organoid sections. Arrows indicate HOPX+/ZsGreen+ progenitor regions. Astrisks indiate HOPX-/ZsGreen-progenitor regions. X indicates HOPX labeling artifact. Scale bars: (A) 5 weeks = 100um, (A) 10/20 weeks = 200um, (B) = 100um, (C) = 50um.

To determine whether the CTNNB1 p.Gln76* variant alters the neuronal cell types produced in cortical organoids, the proportion of various progenitor and neuronal groups were compared between genotypes without consideration of the COR-LT system. At week 10, the proportion of deep layer excitatory neurons in CTNNB1 WT/Q76* organoids was reduced compared to CTNNB1 WT/WT organoids (Fig. 4A-D, scRNA-seq = 23.6% vs 36.1%; IHC = 14.8%, +/− 2.0% vs 21%, +/− 1.3%, p = 0.0102). However, the neuronal reduction in CTNNB1 WT/Q76* organoids observed at week 10 was normalized by week 20, as the proportion of both deep layer neurons (CTNNB1 WT/WT = 12%; CTNNB1 WT/Q76* = 12.2%) and upper layer neurons (CTNNB1 WT/WT = 22%; CTNNB1 WT/Q76* = 25.1%) were similar between different organoid genotypes (Fig. 4E-G). This suggests that the CTNNB1 p.Gln76* variant led to delayed neuron production at earlier stages of organoid development which was later normalized as organoid development progressed.

**Figure 4.**
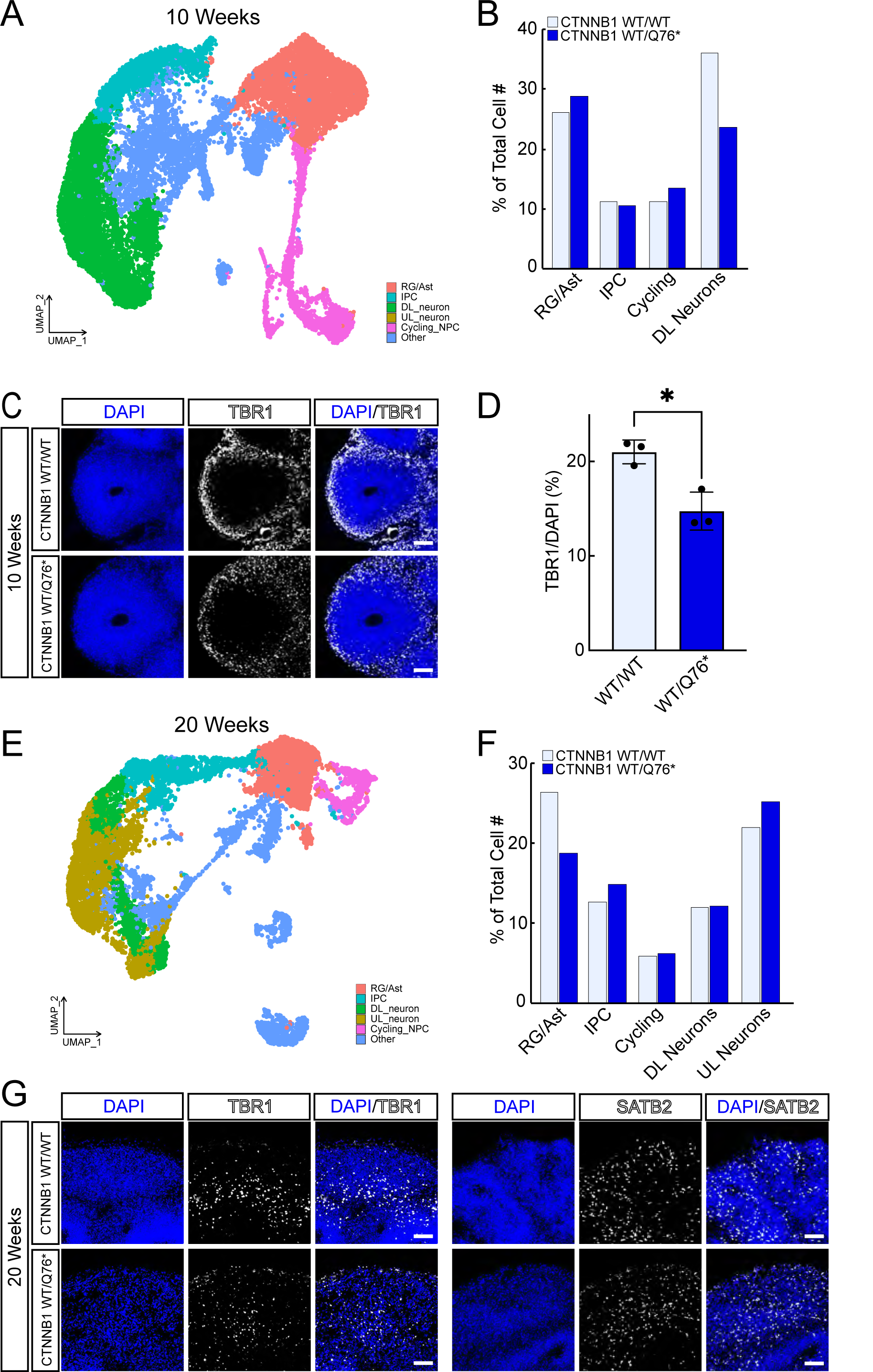
Detection of delayed deep layer neuron production in CTNNB1 WT/Q76* organoids without the COR-LT system. (A) Integrated UMAP of scRNA-seq experiments from 10-week-old ASD organoids with general cell classifications. (B) Proportion of specific cell types in CTNNB1 WT/WT (light blue) and CTNNB1 WT/Q76* (dark blue) 10-week-old organoids as determined by scRNA-seq. (C) IHC detection of TBR1 (white) in 10-week-old ASD organoid sections. (D) Quantification of IHC for TBR1 proportion in individual 10-week-old organoid sections. Error bars = SD, *p<0.05. (E) Integrated UMAP of scRNA-seq experiments from 20-week-old ASD organoids with general cell classifications. (F) Proportion of specific cell types in CTNNB1 WT/WT (light blue) and CTNNB1 WT/Q76* (dark blue) 20-week-old organoids as determined by scRNA-seq. (G) IHC detection of TBR1 (white, left) and SATB2 (white, right), in 20-week-old ASD organoid sections. Scale bars = 100um.

### COR-LT system identification of lineage-specific differences in deep and upper layer neuronal production in CTNNB1 WT/Q76* cortical organoids

We then determined whether use of the COR-LT system could generate additional insight into the differences between these isogenic CTNNB1 WT/WT and CTNNB1 WT/Q76* organoids. scRNA-seq uncovered all four lineages of neurogenesis in both CTNNB1 WT/WT and CTNNB1 WT/Q76* organoids: vRG direct (“no color”, tdTomato-/ZsGreen-); vRG->IPC indirect (“red”, tdTomato+/ZsGreen-); oRG direct (“green”, tdTomato-/ZsGreen+); and oRG->IPC indirect (“yellow”, tdTomato+/ZsGreen+) neurogenesis. Similar to what was found in control organoids, the vast majority of neurons in both *CTNNB1* genotypes at both time points expressed tdTomato, indicating they were generated indirectly through IPCs (Fig. 5A-E). Interestingly, the CTNNB1 p.Gln76* variant led to the reduction of deep layer neurons derived from the “red” lineage at 10 weeks (CTNNB1 WT/WT = 88.2%; CTNNB1 WT/Q76* = 63.9%), and a corresponding increase in the production of neurons derived from “yellow” lineage (CTNNB1 WT/WT = 6.6%; CTNNB1 WT/Q76* = 29.9%) (Fig. 5E).

**Figure 5.**
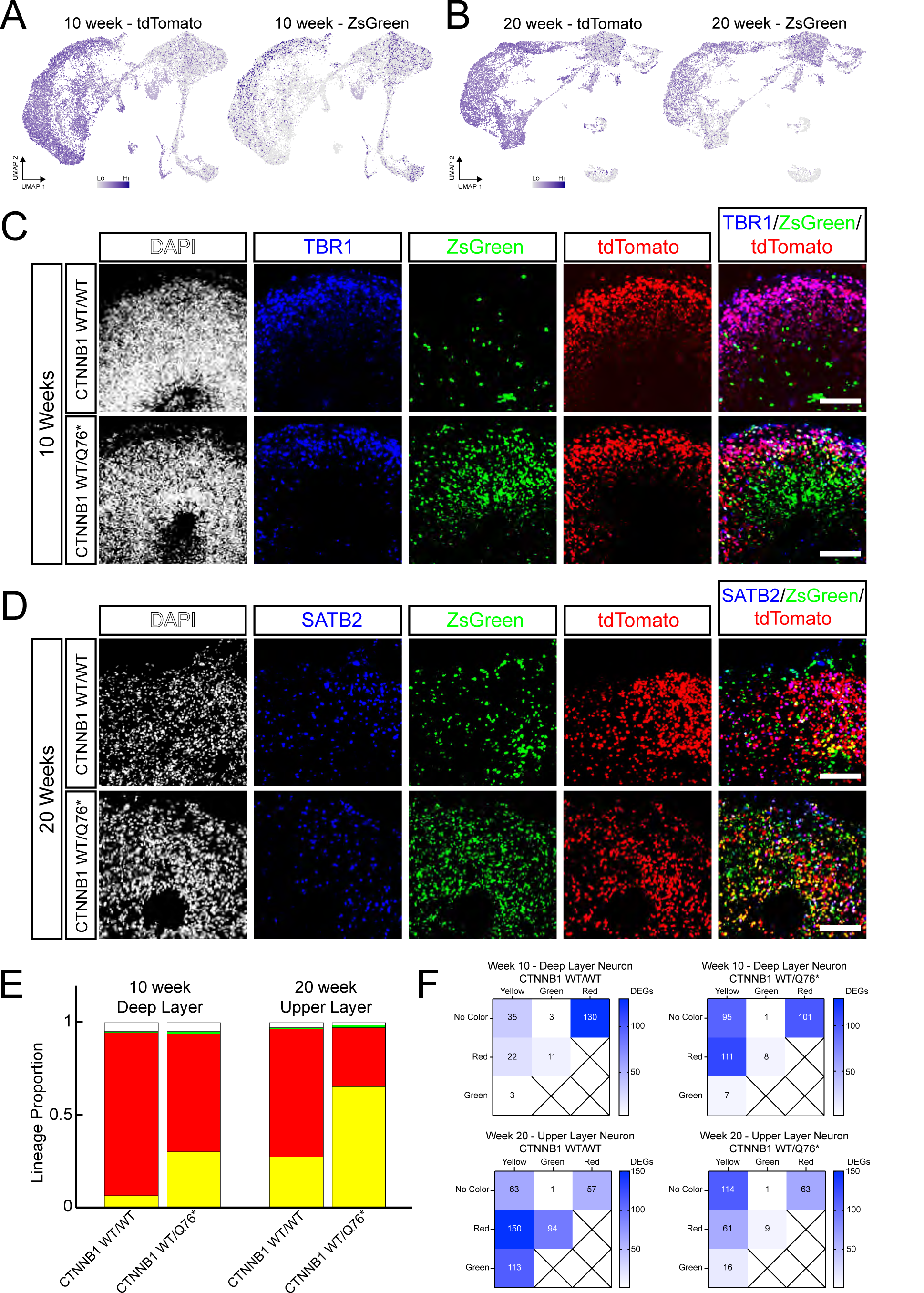
Detection of neuronal production through different progenitor cell lineages in CTNNB1 WT/Q76* organoids with the COR-LT system. (A, B) UMAPs of tdTomato and ZsGreen expression in scRNA-seq experiments from 10-week-old (A) and 20-week-old (B) ASD organoids. (C) IHC detection of TBR1 (blue) with ZsGreen (green) and tdTomato (red) expression in 10-week-old ASD organoid sections. (D) IHC detection of SATB2 (blue) with ZsGreen (green) and tdTomato (red) expression in 20-week-old ASD organoid sections. (E) Lineage proportions of deep layer neurons in 10-week-old ASD organoids (left) and upper layer neurons in 20-week-old ASD organoids (right) as determined by reporter expression in scRNA-seq data. (F) Matrices indicating the number of differentially expressed genes (p_val_adj < 0.05) between pair-wise comparisons of different lineage neurons. Scale bars = 100um.

This suggests that oRGs were involved in neuronal production at earlier stages in CTNNB1 WT/Q76* organoids compared to CTNNB1 WT/WT organoids. As noted above, ZsGreen was observed in CTNNB1 WT/Q76* organoids, but not in CTNNB1 WT/WT organoids at 5 weeks (Fig. 3A, B), suggesting that oRGs were produced earlier in organoids with this variant. IHC of 5-week-old organoid sections revealed that HOPX and corresponding ZsGreen expression was indeed present in CTNNB1 WT/Q76* organoids but absent in CTNNB1 WT/WT organoids (Fig. 3C), while, by 10 weeks, HOPX expression was detected in progenitor regions of both groups (Fig. 3C). The number of deep layer neurons expressing ZsGreen was higher in CTNNB1 WT/Q76* organoids compared to CTNNB1 WT/WT organoids at 10 weeks, likely because of the extended oRG production period in CTNNB1 WT/Q76* organoids (Fig. 5E – “green” + “yellow”; CTNNB1 WT/WT = 7.0%; CTNNB1 WT/Q76* = 31.5%). By 20 weeks, the proportion of upper layer neurons generated via the oRG->IPC lineage was substantially increased in CTNNB1 WT/Q76* organoids (Fig. 5E - “yellow”; CTNNB1 WT/WT = 27.2%; CTNNB1 WT/Q76* = 65.1%), while the vRG->IPC lineage predominated in CTNNB1 WT/WT organoids (Fig. 5E – “red”, CTNNB1 WT/WT = 69.3%, CTNNB1 WT/Q76* = 32.1%). Thus, although the overall proportion of upper layer neurons was relatively similar between CTNNB1 WT/WT and CTNNB1 WT/Q76* organoids at 20 weeks (Fig. 4), the lineages of these neurons were substantially altered due to the CTNNB1 p.Gln76* mutation.

### Gene expression profiles of deep and upper layer neurons of different lineage in isogenic ASD organoids were distinct and altered by the CTNNB1 p.Gln76* variant

If neurons generated through different lineages are transcriptionally distinct, then alterations in lineage-specific neuronal production might lead to functional changes in the brain, even if the overall proportion of neuronal subtypes are maintained. Therefore, transcriptional differences within the same neuronal subtypes but of different progenitor lineage were determined in CTNNB1 WT/WT and CTNNB1 WT/Q76* organoids via scRNA-seq. For these comparisons, we focused on deep layer neurons from 10-week-old organoids and upper layer neurons from 20-week-old organoids, as these neuronal subtypes made up the majority of neurons within organoids at their respective time points.

In 10-week-old organoids, most deep layer neurons were derived through one of two major lineages: vRG->IPC (“red”), or oRG->IPC (“yellow”). A total of 22 DEGs were identified when comparing “yellow” vs “red” deep layer neurons in CTNNB1 WT/WT organoids, with 3 SFARI ASD risk genes differentially expressed (Fig. 5F, Supplemental Table 6, https://github.com/lbury3/cor_lt/). Similarly, a total of 111 differentially expressed genes (DEGs) in week 10 CTNNB1 WT/Q76* organoids were found between “yellow” vs “red” deep layer neurons, 19 of which are SFARI ASD risk genes (Fig. 5F, Supplemental Table 6, https://github.com/lbury3/cor_lt/). Additional pairwise comparisons between deep layer neurons of different lineages were also performed (“red” vs “no color”; “green” vs “no color”; “yellow” vs “no color”; “yellow” vs “red”; “yellow” vs “green”; “green” vs “red”). DEGs were found in all comparisons in both CTNNB1 WT/WT and CTNNB1 WT/Q76* organoids (Fig. 5F). However, as seen in the “yellow” vs “red” lineage comparison, the number of DEGs in the same pairwise comparison but different organoid genotype varied considerably (Fig. 5F). Moreover, the specific genes that were differentially expressed in an individual pairwise comparison differed between organoids of different genotype, including a number of SFARI ASD risk genes (Supplemental Table 6, https://github.com/lbury3/cor_lt/).

Most upper layer neurons in 20-week-old organoids were derived from either the vRG->IPC (“red”) lineage or the oRG->IPC (“yellow”) lineage as well. A total of 150 DEGs were identified between “red” and “yellow” lineage upper layer neurons from CTNNB1 WT/WT 20-week-old organoids, compared to 61 DEGs in the “red” vs “yellow” lineage comparison in CTNNB1 WT/Q76* organoids (Fig. 5F, Supplemental Table 7, https://github.com/lbury3/cor_lt/). Additional pairwise comparisons between neuronal groups of distinct lineage indicated differences in the number of DEGs found in upper layer neurons from CTNNB1 WT/WT organoids compared with upper layer neurons from CTNNB1 WT/Q76* organoids (Fig. 5F). These results indicate that both upper layer and deep layer neurons derived from different lineages possessed distinct gene expression profiles in ASD organoids, as is observed in upper layer neurons from control organoids (Fig. 2G, H, Supplemental Fig. 5C). Additionally, these distinct gene expression profiles were altered depending on *CTNNB1* variant status.

To determine the biological relevance of these differences, pairwise GSEA was performed across different lineages within the same cell line. We then determined whether each lineage possessed a unique set of gene ontology (GO) terms and whether the *CTNNB1* mutation changed the landscape of the GO terms enriched in the “between lineage” comparisons. For deep layer neurons in 10-week-old organoids, a number of distinct GO terms were enriched in various lineages (Fig. 6A, B). The pattern of GO term enrichment was relatively similar in both CTNNB1 WT/WT and CTNNB1 WT/Q76* organoids, with synaptic signaling terms generally enriched in IPC-derived tdTomato+ neurons (vRG->IPC “red” and oRG->IPC “yellow” lineages, Fig. 6A, B), similar to what was found in upper layer neurons in control organoids at 4 months (Supplemental Fig. 5C). There were some differences between genotypes, however, such as the enrichment of genes linked with “regulation of synaptic plasticity” in “yellow” lineage neurons compared to “red” lineage neurons in CTNNB1 WT/Q76* organoids which was not found in the “yellow” vs “red” neuronal lineage comparison in CTNNB1 WT/WT organoids (Fig. 6A, B).

**Figure 6.**
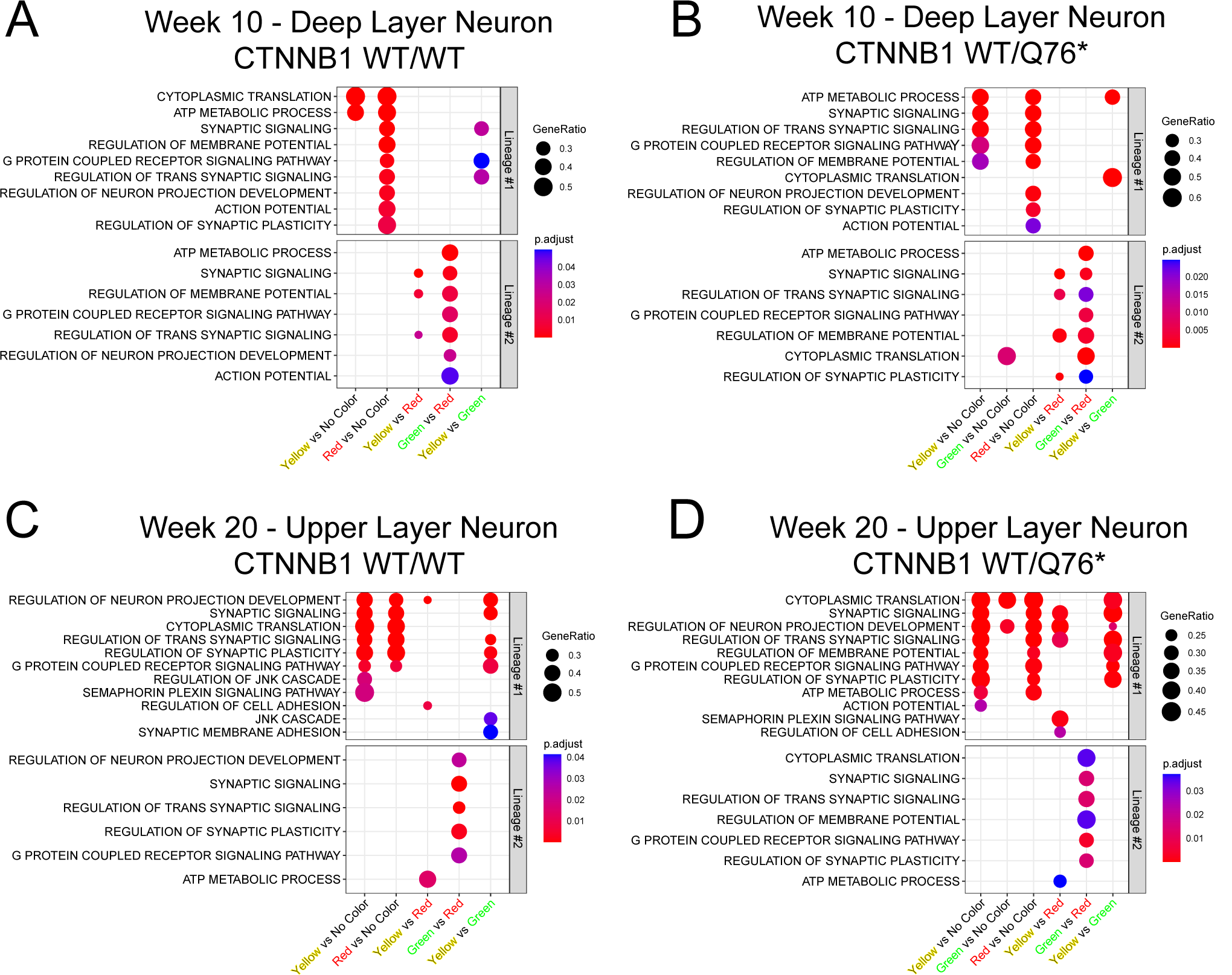
Differential biological pathway enrichment in neurons of distinct progenitor cell lineage within individual ASD cell lines. (A, B) Enrichment of GO terms determined by pairwise GSEA comparisons of different progenitor cell lineages of deep layer neurons from 10-week-old CTNNB1 WT/WT (A) or CTNNB1 WT/Q76* (B) organoids. Top box (“Lineage #1”) indicates GO terms enriched in neurons of the first lineage listed on the x-axis, bottom box (“Lineage #2”) indicates GO terms enriched in neurons of the second lineage listed on the x-axis. (C, D) Enrichment of GO terms determined by pairwise GSEA comparisons of different progenitor cell lineages of upper layer neurons from 20-week-old CTNNB1 WT/WT (C) or CTNNB1 WT/Q76* (D) organoids. For all GSEA analyses, GO terms were preselected for visualization and p-values were adjusted with the Benjamini-Hochberg correction.

Pairwise GSEA comparisons of upper layer neurons in 20-week-old organoids were also generally similar between genotypes. In both CTNNB1 WT/WT and CTNNB1 WT/Q76*, tdTomato+ IPC-derived neurons from either vRG->IPC “red” or oRG->IPC “yellow” lineages were enriched in GO terms related to synaptic signaling and development, compared to other lineages (Fig. 6 C, D), which was again similar to what was found in neurons from control organoids at 4 months (Supplemental Fig. 5C) and 10-week-old isogenic *CTNNB1* lines (Fig. 6A, B). However, genotype-dependent differences were again noted in the “yellow” vs “red” lineage comparison. Specifically, “synaptic signaling” and “semaphorin plexin signaling pathway” were enriched in “yellow” lineage neurons compared to “red” lineage neurons in CTNNB1 WT/Q76* organoids whereas these terms were not enriched in the “yellow” vs “red” lineage comparison in CTNNB1 WT/WT organoids (Fig. 6C, D). Genotype-dependent enrichment of these neurodevelopmental GO terms in isogenic cortical organoids suggest that biologically relevant processes are altered in a lineage-specific fashion due to the pathogenic CTNNB1 p.Gln76* mutation.

To further distinguish the effects of CTNNB1 p.Gln76* mutation on neuronal identify, the transcriptional profiles of neurons from identical subtype and identical lineage in CTNNB1 WT/WT and CTNNB1 WT/ Q76* organoids were directly compared. In deep layer neurons from 10-week-old organoids, DEGs between CTNNB1 WT/WT and CTNNB1 WT/Q76* were detected in three out of four lineages (Fig. 7A, Supplemental Table 8, https://github.com/lbury3/cor_lt/; vRG->IPC “red”, oRG->IPC “yellow”, vRG direct “no color”). In upper layer neurons from 20-week-old organoids, DEGs between CTNNB1 WT/WT and CTNNB1 WT/Q76* were detected in two out of four lineages (Fig. 7B, Supplemental Table 9, https://github.com/lbury3/cor_lt/; vRG->IPC “yellow”, oRG->IPC “red”). The different number of DEGs in each lineage indicates that the *CTNNB1* variant altered gene expression profiles differently in distinct lineages.

**Figure 7.**
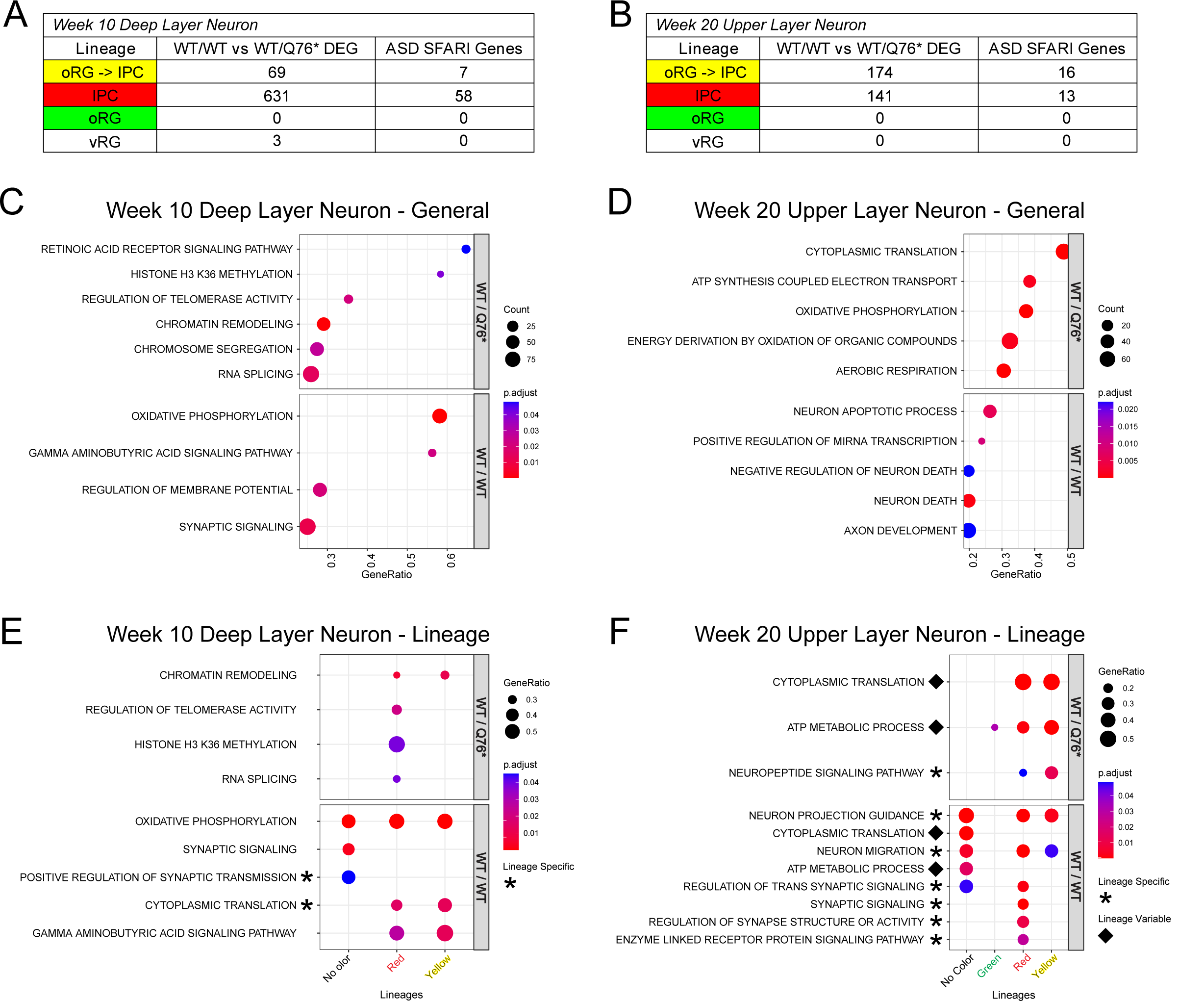
Differential biological pathway enrichment in neurons of distinct progenitor cell lineage between individual ASD cell lines. (A, B) Tables indicating the number of differentially expressed genes (p_val_adj < 0.05) between deep layer (A) or upper layer (B) neurons of the same progenitor cell lineage but different genotypes in 10-week-old (A) or 20-week-old (B) organoids, respectively. (C) Differentially enriched GO terms determined by GSEA comparisons of all deep layer neurons from 10-week-old CTNNB1 WT/WT and CTNNB1 WT/Q76* organoids. (D) Differentially enriched GO terms determined by GSEA comparisons of all upper layer neurons from 20-week-old CTNNB1 WT/WT and CTNNB1 WT/Q76* organoids. (E) Differentially enriched GO terms determined by GSEA comparisons of deep layer neurons from 10-week-old organoids of the same lineage but different genotype. (F) Differentially enriched GO terms determined by GSEA comparisons of upper layer neurons from 20-week-old organoids of the same lineage but different genotype. Asterisks indicate differentially enriched GO terms between genotypes that are found in specific lineages but not general neuronal comparisons. Diamonds indicate GO terms that are differentially enriched in neurons of different lineages depending on genotype. For all GSEA analyses, GO terms were preselected for visualization and p-values were adjusted with the Benjamini-Hochberg correction.

Lineage-specific differences between CTNNB1 WT/WT and CTNNB1 WT/Q76* neurons could be due to the effect of the p.Gln76* mutation on distinct lineage-specific neuronal populations, or due to more general differences between CTNNB1 WT/WT and CTNNB1 WT/Q76* neurons, regardless of lineage identity. To distinguish between these possibilities, GSEA analysis comparing CTNNB1 WT/WT vs CTNNB1 WT/Q76* deep layer neurons in 10-week-old organoids (Fig. 7C) and upper layer neurons in 20-week-old organoids (Fig. 7D) was performed without regard for lineage distinctions identified by the COR-LT system. These results were compared with lineage-specific GSEA comparisons between neurons from CTNNB1 WT/WT organoids and neurons from CTNNB1 WT/Q76* organoids (Fig. 7E, F). While many GO terms overlapped between the general and lineage-specific GSEA, a number of GO terms were identified only in lineage-specific groups (Fig. 7E, F) and not in the general comparisons between different cell lines if the COR-LT lineage labeling information was not considered (Fig. 7C, D). For example, in deep layer neurons from 10-week-old organoids, genes involved in “positive regulation of synaptic transmission” were enriched in CTNNB1 WT/WT vRG-direct “no color” neurons while genes involved in “cytoplasmic translation” were enriched in CTNNB1 WT/WT vRG->IPC “red” and oRG->IPC “yellow” lineage neurons (Fig. 7E). In upper layer neurons from 20-week-old organoids, genes involved in the “neuropeptide signaling pathway” were enriched in CTNNB1 WT/Q76* vRG->IPC “red” and oRG->IPC “yellow” lineage neurons, while multiple GO terms related to synaptic signaling and neural development that were not found in the general GSEA comparison were found in CTNNB1 WT/WT neurons in a variety of individual lineages (Fig. 7F). Furthermore, GO terms that overlapped between lineage-specific GSEA and general GSEA were found to be enriched in some neuronal lineages and not others. For example, for deep layer neurons in general, “regulation of telomerase activity”, “histone H3 K36 methylation”, and “RNA splicing” were all upregulated in CTNNB1 WT/Q76* organoids (Fig. 7C). Lineage-specific GSEA confirmed that these terms were upregulated in some deep layer neurons from CTNNB1 WT/Q76* organoids, but only those derived from the vRG->IPC “red” lineage (Fig. 7E).

For GSEA analyses of isogenic ASD cell lines, GO terms presented in Figures 6 and 7 were preselected for visualization. Full lists of significantly enriched GO terms for all comparisons are presented in Supplemental Tables 10 and 11 (https://github.com/lbury3/cor_lt/).

These findings in both control and isogenic ASD organoids strongly support the notion that neurons of the same neuronal subtype with different progenitor lineages possessed important transcriptional, and likely functional, differences. In addition, the ASD-linked CTNNB1 p.Gln76* variant altered these lineage-specific differences.

### Identification of common markers for distinct lineages

To determine whether a common gene expression profile across control and isogenic ASD cell lines could be established for neurons of different lineages, DEGs between vRG->IPC “red” and oRG->IPC “yellow” lineage upper layer neurons were compared between control (4 month), CTNNB1 WT/WT (20 week), and CTNNB1 WT/Q76* (20 week) genotypes (Supplemental Fig. 9A, B). Comparisons between other lineages were excluded from this analysis due to the small number of cells in the “green” and “no color” lineages.

Most DEGs were unique to each genotype. However, a single gene was enriched in both “red” and “yellow” lineage neurons from all three cell lines - *CCNI* for oRG->IPC “yellow” lineage neurons and *LINC00632* for vRG->IPC “red” lineage neurons (Supplemental Fig. 9A, B). *CCNI* is present in post-mitotic neurons and codes for the cyclin I protein, which acts with CDK5 to regulate neuronal apoptosis,^38^ while *LINC00632* is not well characterized in the brain. Interestingly, the gene *EPHA3* is significantly enriched in the oRG->IPC “yellow” neuronal lineage of both control and CTNNB1 WT/WT neurons (Supplemental Fig. 9A). EPHA3 is part of the Eph receptor tyrosine kinase family, which bind membrane-associated ephrin ligands and play important roles in neuronal migration and axonal targeting during brain development.^39^ In both mouse and macaque, *EPHA3* localization and timing correlates with neurogenesis through oRGs in each respective species.^20, 21, 40–44^ Enrichment of *EPHA3* in oRG-derived ZsGreen+ neurons in organoids from two different genetic backgrounds suggests that this gene might be a general marker for oRG-derived neurons.

## Discussion

In this study we developed the COR-LT system for determining the progenitor cell lineage of neurons in human cortical organoids. Using this genetic labeling system, the progenitor cell lineage of neurons can be identified via the expression of specific fluorescent reporters that were initially activated in different NPC subtypes: *EOMES* for IPCs and *HOPX* for oRGs. We confirmed that this system faithfully determined the neuronal progenitor lineage of neurons in cortical organoids, which was used to uncover several important and heretofore underappreciated avenues through which neuronal diversity is guided by neuronal progenitor lineage.

### Indirect neurogenesis is the predominant form of neuronal production in human cortical organoids

Numerous mouse studies utilizing both live time-lapse imaging and genetic lineage tracing systems demonstrated that a combination of indirect neurogenesis through IPCs and direct neurogenesis from vRGs (and oRGs, rarely) contributes to cortical neuron production.^11, 12, 14, 17, 19–21, 45^ Much less is known about the proportion of neurons produced through indirect vs. direct neurogenesis in humans or human model systems, and the data available has resulted in conflicting interpretations.

Viral lineage tracing suggested that oRGs produce neurons indirectly through IPCs in human fetal brain while macaque oRGs produce neurons both directly and indirectly.^8, 44^ This discrepancy might reflect differences in biology between human and macaque or could be due to differences in whether to classify oRGs and IPCs through morphology or EOMES expression.^8, 44, 46^ Computational lineage tracing in human organoids and human fetal brain has yielded discordant results with some studies suggesting primarily indirect neurogenesis,^47–49^ while others proposing a mixture of direct and indirect.^50, 51^ Conflicting results in these cases might be due to differences in the specific methods used or possibly from the inclusion of stressed cells in lineage analysis.^49^ Genetic barcoding approaches in organoids have been used to trace the lineage of individual cells back to an initial founder cell,^52–54^ but lineage information from specific progenitor subtype populations has not been established. To our knowledge, lineage tracing from an entire genetically defined progenitor population in a human system has not been performed.

We have employed lineage tracing by tagging all neurons that were produced through progenitors expressing the canonical IPC marker *EOMES*, and have found that the vast majority of neurons in both control and ASD isogenic organoids were produced indirectly through IPCs (Fig. 2, Fig. 5). This result is somewhat surprising, based on studies in other mammals, especially mice. However, reliance on indirect neurogenesis for neuronal production would theoretically lead to a greater number of neurons produced in the human cerebral cortex overall, as primate IPCs can undergo multiple rounds of amplifying divisions before terminally differentiating into neurons.^8, 44^ The expansion of the human cortex compared to other species is thought to be driven, at least in part, by an increase in neuronal production from oRGs in the outer subventricular zone of the developing brain.^4, 55, 56^ An increase of indirect neurogenesis through IPCs might serve as a complementary mechanism to increase neuronal number in the human cortex,^57^ if what is found in cortical organoids reflects neurogenesis in fetal brain development.

### Neuronal progenitor origin leads to lineage-dependent neuronal diversity

In mice, neurons generated indirectly from IPCs are functionally, morphologically, and transcriptionally distinct^17–19^ from directly-generated vRG-derived neurons. Whether this is also the case for human neurons is unknown. Additionally, it is not known whether neurons derived from oRGs are distinct from non-oRG-derived neurons in any biological system. In this study, we used the COR-LT system in combination with scRNA-seq to identify transcriptomic differences between neurons from different lineages, including in oRG-derived neurons. These experiments revealed that many of the genes that distinguish these different lineages play important roles in brain development and disease. Further investigation of lineage-specific neuronal differentiation will lead to increased understanding of the factors underlying neuronal diversity in the human cerebral cortex.

### Isogenic iPSC lines reveal CTNNB1-dependent lineage specific developmental defects and mechanisms

The COR-LT system can also be utilized to detect subtle alterations in neurogenesis, such as those that occur through different lineages due to a disease variant or mutation. We found that organoids generated from isogenic ASD cell lines were generally similar, with a decrease in neuronal production at 10 weeks that was normalized by week 20 (Fig. 4). We also observed the early production of oRGs due to the ASD-patient-relevant *CTNNB1* variant. When we employed the COR-LT system, we detected a major shift from vRG->IPC to oRG->IPC neurogenesis due to this variant (Fig. 5), suggesting that neurogenesis through oRGs is potentially an important disease relevant phenotype.

In both control and isogenic ASD organoids, neurons derived through different lineages were found to be transcriptionally distinct (Fig. 2, Fig. 5, Fig. 6, Supplemental Fig. 5). In addition, the ASD-linked *CTNNB1* variant itself was associated with unique transcriptional changes in neurons from various lineages (Fig. 6, Fig. 7). Thus, although the proportion of upper layer and deep layer neurons in each isogenic organoid type were similar, the overproduction of oRG-derived neurons that were transcriptionally distinct could lead to functional differences in these neurons. For example, we found that genes related to the semaphorin/plexin signaling pathway were upregulated in “yellow” lineage upper layer neurons compared to “red” lineage upper layer neurons, but only in organoids with the *CTNNB1* ASD variant (Fig. 6C, D).

If neurons from different lineages are transcriptionally distinct in the human brain, as they are in human organoids, it suggests a potential pathological cellular mechanism for neurological disorders where genetic studies have linked aberrant fetal cortical development with the disease. In many of these diseases, gross morphological changes in patient brain structure are subtle and pathological mechanisms are unclear. For example, ASD risk gene expression is enriched during mid-fetal cortical development during the height of neurogenesis,^58–60^ and a subset of individuals with ASD also have co-morbid macrocephaly.^25, 33–35, 37, 61^ However, individuals with ASD typically possess morphologically normal brains.^62^ Altering the proportion of neurons derived from distinct lineages could alter cortical function without detectable changes in general neuronal subtypes (e.g. upper layer/deep layer neurons), leading to disease in people with otherwise normal brain anatomy. Lineage-specific genes might therefore be interesting therapeutic targets for modulating the activity of over- or under-produced neuronal lineages.

### Study limitations

*HOPX* possesses three major isoforms, A, B, and C.^63^ To activate the COR-LT system in oRGs, Dre recombinase was integrated into the C-terminus of *HOPX* isoform C, which is upstream of the C-termini of isoforms A and B.^63^ Since isoforms A and B are out-of-frame compared to isoform C, integration of Dre recombinase at the chosen site leads to the replacement of the last 23 amino acids of isoforms A and B with 7 alternative amino acids (LRAEEVF). The functional consequences of this truncation in a subset of *HOPX* isoforms from a single allele are unclear. Altered isoform expression might lead to gene expression changes in oRG cells themselves. However, *HOPX* is not expressed in neurons (Fig. 1, Supplemental Fig. 8) making it less likely that alterations in *HOPX* causes the differential gene expression found in the various lineage comparisons performed in this study. Importantly, the WT *HOPX* alleles in both control and ASD lines from our study were unperturbed and robust expression of HOPX protein was detected in organoids from all cell lines via IHC (Fig. 1, Fig. 3). Additionally, heterozygous knockout mice for *HOPX* are developmentally and behaviorally normal with normal brain anatomy.^64, 65^

Because of this frame shift, Dre recombinase protein will not be produced from isoform A or isoform B mRNA. *HOPX* isoform C is expressed in human oRGs during fetal development (Arnold Kriegstein, personal communication) and is expressed in the organoids in this study. However, if either isoform A and B are expressed while isoform C is not expressed in an individual oRG cell, the COR-LT system would not be activated, leading to a potential undercount of oRG-derived neurons. Furthermore, although co-localization of ZsGreen with the TBR1 neuronal marker in “pre-activated” organoids was high, there was a small population of neurons that did not express ZsGreen (Supplemental Fig. 1B, C). While the exact mechanism of producing these ZsGreen-negative neurons is unclear, if the ZsGreen reporter is silenced or not initially activated in some neurons, it would also lead to an undercount of oRG-derived neurons. We do not believe that these limitations affect our interpretations. For example, in spite of these potential caveats, we were still able to detect a significant number of neurons that were from an oRG-derived lineage as well as transcriptional differences between oRG-derived and non-oRG-derived neurons in organoids from multiple cell lines, including the isogenic CTNNB1 lines.

Sanger sequencing of the WT *EOMES* allele (without COR-LT construct integration) detected a 13bp deletion within the last exon of the gene for the CTNNB1 WT/WT and CTNNB1 WT/Q76* cell lines. This deletion caused a frame-shift which led to the final 26 amino acids to be replaced with a novel 26 amino acid sequence at the C-terminus. While this amino acid change does not affect the critical T-Box domain through which EOMES binds DNA, the alteration might affect some functions of the gene. However, the function of EOMES from the edited allele could compensate for any reduced activity from the unintentionally mutated allele. Sanger sequencing of the edited allele indicated proper integration of Cre recombinase, which yields a full-length copy of EOMES with the short 2A peptide attached to the C-terminal. Notably, humans and mice with heterozygous *EOMES* mutations are healthy^66, 67^ and EOMES protein is readily detected in organoids from both cell lines (Fig. 1, Supplemental Fig. 7). Additionally, the large number of tdTomato+ neurons in CTNNB1 WT/WT and CTNNB1 WT/Q76* organoids indicates that neurons are abundantly produced through *EOMES*+ IPCs. Also, differential gene expression comparisons were only performed between isogenic cell lines, which have the same genetic background, including this mutation in *EOMES*. Therefore, it is unlikely that the mutation in the unedited *EOMES* allele would affect the general conclusions of this study.

Finally, our studies were performed in cortical organoids, not in the intact, developing human cortex, so it is unclear whether our findings also apply to the human brain. Organoids lack a number of key cell types and molecular signaling sources that play important roles in brain development, such as microglia, connectivity with other brain regions, exposure to cerebrospinal fluid, and a functioning vasculature system. It remains to be determined how these important contributors to cortical development would affect the results presented here. However, a recent large-scale single cell atlas study has indicated that, even without these additional components, both progenitor cells and neurons in cortical organoids faithfully recapitulate the transcriptomic profiles observed in fetal cortical development.^51^ Therefore, we believe that our conclusions drawn from organoids most likely reflect basic developmental processes in the human cerebral cortex, but further studies are needed to confirm this notion.

### Future directions

The COR-LT system can theoretically be implemented in any human cell line, and, by varying the gene targets for insertion of Cre and Dre recombinases, can be used to assess neurogenesis from any neural progenitor population. For example, insertion of the recombinases into ATOH1 and KIRREL2, followed by production of cerebellar organoids, would allow the assessment of granule and Purkinje cell neurogenesis, respectively. The effect that different neurological diseases have on neurogenesis in cortical organoids could also be tested by installing the system into various disease cell lines. The development of isogenic cell lines testing the effect of disease-relevant mutations could be readily generated by performing a single genome-editing step in a cell line with the COR-LT system previously installed, as demonstrated by the isogenic ASD cell lines described in this study. Evaluating neurogenesis in additional control cell lines with this system could potentially provide more statistical power for determining lineage-specific gene expression differences in neurotypical development. In addition to human development, evolutionary questions might also be addressed by installing the COR-LT system (or functionally similar system) into non-human cell lines and comparing neuronal lineages between organoids from different species.

Because the goal of this study was to assess population-level alterations in neuronal lineage and to identify sufficient neurons from each lineage to assess gene expression differences, clonal analysis of individual progenitor cells was not attempted. Modified versions of the COR-LT system might address the neurogenic potential of individual progenitor cells in human cortical organoids. An inducible COR-LT system would also be useful in determining the time-course of neurogenesis from various progenitor lineages.

In conclusion, we demonstrate that the COR-LT system is a reliable tool for determining the progenitor cell lineage of neurons in cortical organoids. Further implementation of this system should yield valuable insights into cortical development in both health and disease, as well as development of any other human brain region in which organoids can be produced.

## Acknowledgements

We would like to thank Paul Tesar, Ashleigh Schaffer, Helen Miranda, Fulai Jin, and Marissa Scavuzzo for their helpful critiques of the manuscript. We would also like to thank Jaejin Eum for assistance with immunohistochemistry analysis as well as Konstantin Leskov and Marissa Scavuzzo for assistance with single-cell sequencing analysis. This work was supported by grants from the NIH and NIMH (R01 MH113106 and MH114601) to A.W.-B. and by grants from the Hartwell Foundation (Hartwell Biomedical Research Fellowship), the Brain and Behavior Research Foundation (NARSAD Young Investigator Grant, ID 26078), and the NIH (5 TL1 TR 0002549-03) to L.A.D.B. Imaging was performed on microscopes acquired and maintained by the CWRU SOM Light Microscopy Core Facility. Grants supporting equipment utilized in this publication include NIH Grant S10-OD016164 (SP8), NIH Grant S10-RR021228 (DMI6000, DM6000), and NIH Grant ORIP S10OD024981 (Nanozoomer). Training and assistance was graciously provided by Patricia Conrad, Richard Lee, Kelley Carr, and Philip Ropelewski, all of the CWRU SOM Light Microscopy Core.

## Author Contributions

Conceptualization, L.A.D.B., S.F., A.W.-B.; Methodology, L.A.D.B., S.F., A.W.-B; Formal Analysis, L.A.D.B., S.F.; Investigation, L.A.D.B, S.F.; Writing – Original Draft, L.A.D.B., S.F.; Writing – Review and Editing, L.A.D.B., S.F., A. W.-B.; Visualization, L.A.D.B., S.F.; Supervision, L.A.D.B., A.W.-B.; Funding Acquisition, L.A.D.B., A.W.-B.

## Methods

### iPS cell lines

Studies utilizing the cell lines described in this paper were approved by the Institutional Review Board of Case Western Reserve University and were previously described.^25^ The cell line “Clay” was utilized as the control cell line in this study and “Arch” was utilized as the ASD cell line in this study. The Arch cell line possessed the *CTNNB1* Q76* variant. The *CTNNB1* Q76* variant was corrected to the WT variant via CRISPR/Cas9 gene editing as described in ^68^ and below and verified via MiSeq.^68^ All cell lines were grown on vitronectin (Thermo Fisher) coated plates and maintained in mTeSR+ media (STEMCELL Technologies). For routine culture, cells were passaged every 5-7 days or once cells reached 70-80% confluency. Regular testing for mycoplasma was consistently negative for all cell lines.

### Plasmid construction for COR-LT system and *CTNNB1* Q76* correction

All plasmids used in these studies are available from Addgene. For all primers referenced below, please see Supplemental Table 12 (https://github.com/lbury3/cor_lt/) for sequence. All primers in this study were obtained from Integrated DNA Technology, Morrisville, NC.

#### Generation of stop-rox ZsGreen reporter construct targeting H11 locus

The original Zsgreen1 fragment within Addgene #51274 was dropped via BamHI and SacI double digestion. Addgene #51274 template with primers GFP-BamHI-S and ZsGFP-NLS-FLG-EcoRI-SacI-R were used to generate a modified Zsgreen1 fragment so that the NLS sequence, FLG tag and additional EcoRI restriction site was added to the Zsgreen1 C-terminus. The modified Zsgreen1 fragment was then ligated to the previously double digested Addgene #51274 vector (both insert fragment and vector were digested with BamI and SacI), generating the CAG-roxSTOProx-Zsgreen-NLS-FLG construct. To add homologous targeting arms to this construct, Addgene #51546 was digested with MreI and EcoRI to drop a 1166bp DNA fragment while the 7676bp H11 vector was saved for future use. CAG-roxSTOProx-Zsgreen-NLS-FLG was also digested with MreI and EcoRI. The insert (2,063bp) from this digestion was saved and used for insertion into the 7676bp H11 vector. This final construct was named H11-CAG-roxSTOProx-Zsgreen-NLS.

#### Generation of stop-flox tdTomato reporter construct targeting AAVS1 locus

The loxpSTOPlox (LSL) fragment from Addgene #89587 was amplified with primers CAG-LSL-MreI-F and CAG-LSL-EcoRV-R. Addgene #89587 was then digested with MreI and EcoRV, and the 7701bp digested vector was saved for future use. The previously generated LSL fragment was ligated to this digested 7701bp vector which created the AAV_LSL plasmid. The NLS-tdTomato fragment was then amplified using Addgene #83029 with primers AAV_LSL_nls_Tdtomato_GS_F and AAV_LSL_nls_Tdtomato_GS_R for future Gibson Assembly. The AAV_LSL plasmid was linearized with EcoRV, also for future Gibson Assembly. Gibson Assembly was then performed using the NLS-tdTomato fragment and linearized AAV_LSL plasmid, which created the final AAVS1_LSL_NLS_tdTomato plasmid.

#### Construction of T2A-NLS-Cre targeting EOMES C-terminus

The left homologous arm was amplified using genomic DNA from the control iPSC line Chap^25^ as template with primers EOMES-C-Cre-LHA_fwd and EOMES-C-Cre-LHA_rev. The right homologous arm was amplified by using genomic DNA from the Chap iPSC line with primers EOMES-C-Cre-RHA_fwd and EOMES-C-Cre-RHA_rev. The Cre fragment was then amplified using Addgene #26646 as template with primers T2A-NLS-Cre_fwd and T2A-NLS-Cre_rev. The vector was amplified using pUC19 as template with primers EOMES-C-Cre-UC19_fwd and EOMES-C-Cre-UC19_rev. Finally, Gibson Assembly was performed to assemble the four previously generated fragments which created the final construct, EOMES-2A-Cre.

#### Construction of T2A-NLS-HA-Dre targeting HOPX C-terminus

The left homologous arm was amplified using genomic DNA from a control iPS cell line (“Chap” as described in ^25^) as template with primers 5’HA_HOPX_F1 and 5’HA_HOPX_R1. The right homologous arm was amplified using genomic DNA as in step 1 with primers 3’HA_HOPX_F1 and 3’HA_HOPX_R1. The Dre fragment was amplified using Addgene #51272 as template with primers T2A-NLS-HA-Dre-F and T2A_HA_DRE_pA_R1. The vector was amplified using pUC19 as template with primers pUC19_HOPX_F1 and pUC19_HOPX_R1. Finally, Gibson Assembly was utilized to assemble the four previously generated fragments which created the final construct, HOPX-2A-Dre.

#### Construction of donor plasmid for CTNNB1 Q76* correction

Using genomic DNA from the Arch iPSC line as template, a 1.6kb band was amplified via a pair of primers (CTNNB1-BamHI-F and CTNNB1-KpnI-R) that were approximately 800bp upstream and downstream of *CTNNB1* c. 226C, respectively. The pUC19 plasmid (NEB, Catalog number: N3041A) was then linearized with KpnI and BamHI and the 1.6kb amplified region was inserted into the linearized pUC19 vector. As the generated donor plasmid will be either with the *CTNNB1* mutation (amplified from the mutant allele) or without the *CTNNB1* mutation (amplified from the wild type allele), Sanger sequencing was used to confirm the generated donor plasmid.

#### COR-LT system gRNA Construction

For targeting Cas9n to desired loci, Addgene #74630 was modified by replacing the original CBh promoter with a CAG promoter for more robust expression of Cas9n upon transfection. The original CBh promoter was removed by digesting with KpnI and AgeI (NEB). CAG promoter was amplified from pCAG-roxSTOProx-ZsGreen (Addgene #51274) using PrimeSTAR polymerase (Takera) and cloned into the AIO-Puro backbone. This construct contained two gRNA cassettes, each driven by a human U6 promoter. Unique gRNA sequences were cloned into this construct, depending on the specific experiment. The guide RNAs GTTCCCACTCATACAGGACT and TTCACTTAAGAACAAGTAGC were used to target the *CTNNB1* c. 226C>T locus; GGCGCTGCTTAAACCATTTC and GCAAAGTGGCGGCGCTCAGA were used for targeting *HOPX* C-terminus; guide RNAs ATTAATGTCCTCACACTTTA and TATAGTAAAGACACCTCAAA for targeting *EOMES* C-terminus; Addgene #51554 and #51555 for targeting H11 locus; Addgene #59025 and #59026 for targeting AAVS1 locus. In addition, we also generated constructs containing guide RNAs GAGCCACATTAACCGGCCCT and ACCCCACAGTGGGGCCACTA for targeting the AAVS1 locus, which worked equally well.

### iPS cell transfection and generation of clones

For COR-LT system construct integration, iPS cell lines were passaged into a 24-well plate. The next day, medium was changed to Essential 8 (Thermo Fisher) and cells were transfected using Lipofectamine Stem Reagent (Thermo Fisher) via the recommended protocol. 400ng of donor plasmid DNA plasmid was used along with 50ng of each CRISPR/TALEN component. In order to reduce off-target genetic editing events, the Cas9 D10A nickase was used ^69^ in all CRISPR/Cas9 experiments. Transfection proceeded over the course of approximately 24 hours, at which time, media was changed to mTeSR+. In order to enhance selection of cells transfected with EOMES-2A-Cre or HOPX-2A-Dre plus their associated CRISPR gRNA constructs, puromycin (Thermo Fisher) was added at a concentration of 0.5ug/ml for 1-3 days, starting upon completion of transfection.

Once transfected cells reached 70-90% confluency, they were dissociated into single cells via Accutase (Thermo Fisher) incubation and plated at clonal density on a vitronectin-coated 6-well dish. For some experiments, CloneR supplement (STEMCELL Technologies) was added to enhance single cell viability. For H11-ZsGreen, G418 (Thermo Fisher) was added to media upon single-cell plating in order to select for clones with stable genomic integration. When individual clones were large enough to be transferred to a new plate successfully, clones were manually isolated and transferred to individual wells of vitronectin-coated 48-well plates. After 7-10 days, clones were passaged again into triplicate wells of vitronectin-coated 48-plates. After further cell expansion, DNA from one well for each clone was isolated via QuickExtract DNA Extraction Solution (VWR) to be used for genotyping. For clones with positive integration, an additional well of cells was passaged for further expansion, while the final well was used for freezing and subsequent storage.

### Genotyping

Genotyping was accomplished via standard PCR, utilizing a single primer inside a unique region of integrated DNA and a single primer outside of the homologous arm used for targeting to the specific genomic site. This strategy yields a single PCR band of known length if constructs are integrated into the proper genomic sequence and no band if there is no integration or integration at the wrong genomic site. Primers used for genotyping were: H11-ZsGreen-F, H11-ZsGreen-R, AAVS1-TdTomato-F, AAVS1-TdTomato-R, EOMES-2A-Cre-F, EOMES-2A-Cre-R, HOPX-2A-Dre-F, HOPX-2A-Dre-R (Supplemental Table 12, https://github.com/lbury3/cor_lt/). The PCR reaction mixture for all reactions consisted of 2x GoTaq Green Mastermix (Promega), 400nM primer-F, 400nM primer-R, 12% extracted DNA (see above), water up to recommended volume. Amplification was performed with the following temp/time thermocycler settings: 95/2min, [95/30s, 55/15s, 72/90s]x30, 72/5min, 4/hold.

### Sanger sequencing

Sanger sequencing of both modified and unmodified *EOMES* and *HOPX* alleles was performed to identify potential unintended mutations that occurred through genome editing. After all genome editing was completed, genomic DNA from COR-LT system cell lines was purified via the DNeasy Blood and Tissue DNA extraction kit according to the manufacturer’s instructions (Qiagen). Unmodified alleles were amplified via primers located outside of the homologous arm sequences, utilizing the following primers: EOMES-out-F, EOMES-out-R, HOPX-out-F, HOPX-out-R (Supplemental Table 12, https://github.com/lbury3/cor_lt/). For both reactions, the PCR reaction mixture consisted of 2x Q5 HotStart Mastermix (NEB), 500nM primer-F, 500nM primer-R, 8% genomic DNA, water up to recommended volume. Amplification was performed with the following temp/time thermocycler settings: 93/30s, [98/10s, 63.5/30s, 72/2min]x35, 72/2min, 4/hold.

Modified alleles were amplified with a single reverse primer inside the DNA sequence encoding either Cre or Dre recombinase (for EOMES-2A-Cre and HOPX-2A-Dre alleles, respectively) and a single forward primer outside of the 5’ homologous arm used for targeting to the specific genomic site. With this strategy, the c-terminus of the endogenous gene, the entire 2A sequence, and part of the n-terminus of either Cre or Dre recombinase were amplified. Specific primers used were EOMES-mod-F or EOMES-out-F, EOMES-2A-Cre-R, HOPX-mod-F or HOPX-out-F, HOPX-2A-Dre-R (Supplemental Table 12, https://github.com/lbury3/cor_lt/). The PCR reaction mixture for all reactions consisted of 5x PrimeSTAR Buffer (Takara), 200uM dNTPs (Takara), 250nM primer-F, 250nM primer-R, 4% genomic DNA, 50x PrimeSTAR GXL DNA polymerase (Takara), water up to recommended volume. Amplification was performed with the following temp/time thermocycler settings: [98/10s, 60/15s, 68/2min]x30, 4/hold.

All reactions yielded a single band when run on an agarose gel, except for amplification of the unmodified *EOMES* allele in the ASD iPS cell line, which yielded a band of expected size, plus multiple bands of larger size. For reactions that yielded single bands, the resulting DNA from the PCR reaction was purified via EXO-SAP cleanup (Affymetrix). Individual EXO-SAP reactions consisted of 1.55ul dH2O, 0.15ul Exonuclease I, and 0.3ul shrimp alkaline phosphatase per 10ul of PCR product. This reaction mixture was then placed in a thermocycler with the following temp/time settings: 37/15min, 72/15min, 4/hold. For the PCR reaction of the unmodified *EOMES* allele in the ASD iPS cell line, the expected-size band was extracted from agarose gel via the QIAquick Gel Extraction kit (Qiagen) and re-amplified to yield a single band. This PCR reaction was purified via EXO-SAP cleanup as described above. Clean PCR products for all cell lines were sequenced via Eurofins SimpleSeq service with primers spanning the amplified region (Supplemental Table 12, https://github.com/lbury3/cor_lt/).

For the control COR-LT system cell line, sequencing indicated no unintended modifications or mutations in the sequenced sections, in either modified or unmodified alleles. For the ASD COR-LT system cell line, sequencing of the unmodified *EOMES* allele indicated a 13bp deletion within the last exon (NC_000003.12:g.5564_5576del), which resulted in a frameshift mutation that led to the last 26 c-terminal amino acids being replaced with a novel 26 amino acid sequence (LKSIVKTPQKAWEGIMLFTQLPKELF) before termination at a newly generated stop codon. Sequencing of the unmodified *HOPX* allele in the ASD iPS cell line indicated a single base-pair substitution in the 3’ UTR region (NC_000004.12:g.33538A>G), which is unlikely to affect HOPX production. Sequencing of modified *EOMES* and *HOPX* alleles in the ASD COR-LT system cell line yielded no unexpected alterations.

### Cortical organoid generation

For COR-LT system organoids, iPS cells were imaged via fluorescent microscopy one day prior to organoid generation. The location of colonies that expressed either tdTomato or ZsGreen were marked on the cell culture plate and these colonies were removed from culture immediately prior to each organoid generation experiment. For the organoid experiment where both reporters were activated in the iPS cell stage (Supplemental Fig. 1), iPS cells from the control COR-LT system cell line were transiently co-transfected with Cre and Dre recombinases. Individual iPS cell clones were isolated, and clones with 100% co-activation of both reporters (tdTomato and ZsGreen) were used to generate organoids.

Cortical organoids were generated using modified versions of previously published protocols.^29–31^ For the control COR-LT system cell line, mTeSR+ media was changed on the iPS cells approximately six hours prior to organoid differentiation. At this time, (day 0 – “D0”) iPS cells were gently dissociated into single cells via Accutase incubation. Cells were then centrifuged and resuspended in cortex media. Cortex media consisted of GMEM (Thermo Fisher), 20% KnockOut Serum Replacement (Thermo Fisher), 1x Glutamax (Thermo Fisher), 1x MEM-NEAA (Thermo Fisher), 1mM sodium pyruvate (Thermo Fisher), and 1% pen/strep (Thermo Fisher). The cell suspension was subsequently diluted to 90,000 cells/ml and supplemented with 5uM SB431542 (Millipore Sigma), 3uM IWR-1 (Millipore Sigma), and 20uM Y-27632 (Tocris). 100ul of cell suspension (9,000 cells) was then dispensed into individual wells of a low-attachment v-bottom 96-well plate (S-Bio) and returned to an incubator set at 37 degrees/5%CO_2_. Two days later (D2), cortex media was freshly supplemented with SB431542 (5uM), IWR-1 (3uM), and Y-27632 (20uM) and 100ul was added to each organoid well. Every other day from D6-D16, 100ul of organoid media was removed and replaced with cortex media freshly supplemented with 5uM SB431542 and 3uM IWR-1 (“SI supplement”).

Cortical organoids from ASD COR-LT system cell lines could not be generated with the “SI supplement” protocol. Although organoids initially formed and grew well for the first few days in culture, they began to disintegrate after approximately one week and were not recoverable. Therefore, some of the initial phases of organoid differentiation were modified. To form organoids from ASD COR-LT system lines, mTeSR+ media was changed on the iPS cells one day prior to dissociation. The following day (D0), organoids were generated and maintained as described above, with the following exceptions. From D0-D6, cortex media was supplemented with 1uM A-83 (STEMCELL Technologies), 180nM LDN193189 (STEMCELL Technologies), and 3uM IWR-1 (Millipore Sigma). On D8, 180ul media was removed from each v-bottom well and replaced with cortex media supplemented with 1uM A-83 and 180nM LDN193189. In addition, 10uM Y-27632 (Tocris) was added to the media from D0-D4, instead of 20uM from D0-D2. From D18 onward, organoids from both control and ASD COR-LT system lines were cultured in exactly the same fashion.

On D18, organoids were removed from the 96-well dish and washed 1x with differentiation media A. Differentiation media A consisted of DMEM/F12 (Thermo Fisher), 1x N2 supplement (Thermo Fisher), 1x B27 supplement without vitamin A (Thermo Fisher), 1x Glutamax, 1x MEM-NEAA, 100uM 2-mercaptoethanol (Fisher Scientific), 1% pen/strep, and 0.1% amphotericin B (Thermo Fisher). Organoids were then transferred to low-attachment 6-well dishes (Corning) with 3ml differentiation media A per well. Up to 16 organoids were added per well. However, the number of organoids initially added per well was typically lower than 16 and was further reduced over time as organoids were utilized for immunohistochemistry experiments. Organoids were also transferred to new wells if crowding and/or organoid merging occurred. After organoids were added, the 6-well low-attachment plate was placed on an orbital shaker (Thermo Fisher) in a 37 degree/5%CO_2_ incubator and rotated at 85RPM for the remainder of the experiment. Starting on D20, approximately 2.5ml of organoid media was removed and replaced with 2.5-3ml fresh differentiation media A every other day. From D36 onward, 1% Geltrex (Thermo Fisher) was added to the media. To prevent Geltrex precipitation in the media, cold Geltrex was added fresh to cold, differentiation media A aliquots and allowed to warm at room temperature for no more than 30 minutes before adding to organoids. From D70 onward, differentiation media B was used for organoid culture and related media changes. Differentiation media B consisted of Neurobasal media (Thermo Fisher), B27 supplement, (Thermo Fisher), 1x Glutamax, 1x MEM NEAA, 1% pen/strep, and 0.1% amphotericin B. 1% Geltrex was also added immediately prior to each media change, as described above. In addition, 0.5mM cAMP (STEMCELL Technologies), 0.2mM ascorbic acid (Millipore Sigma), 20ng/ml BDNF (Peprotec, STEMCELL Technologies), and 20ng/ml GDNF (Peprotec) were also added fresh, immediately prior to media change. Organoids were cultured in this supplemented media until utilized for immunohistochemistry or single-cell sequencing experiments, with media changes every other day.

### Organoid dissociation, single-cell RNA library generation and sequencing

Organoids were dissociated into single cells utilizing the Worthington Papain Dissociation System kit, following a previously described protocol (https://protocolexchange.researchsquare.com/article/pex-258/v1, ^31^) with the following modifications. 2-5 organoids were used per cell line per experiment. Organoids from the same cell lines were dissociated together so that cells from all organoids of the same cell line were pooled for individual experiments. For four-month control and 20 week ASD organoids, the organoids were removed from culture medium and washed once with PBS. A clean razor blade was then used to cut individual organoids into approximately four to six pieces. These organoid pieces were then washed 3x with PBS in an attempt to remove dead and/or stressed cells from the interior of the organoid. After the third wash, PBS was removed, papain/DNase solution was added, and the organoid pieces were cut into smaller pieces to start the dissociation process as described in (https://protocolexchange.researchsquare.com/article/pex-258/v1, ^31^). Upon dissociation completion, cells were diluted in PBS with 0.04% BSA (Millipore Sigma) and kept on ice.

Immediately after dissociation was completed, single-cell RNA libraries were generated with the Chromium Next GEM Single Cell 3’ Reagent Kit, v3.1 (10X Genomics) following the manufacturer’s instructions. For each experiment, 10,000 cells were targeted for recovery. Purified libraries were shipped to MedGenome (Forest City, CA) for paired-end sequencing via the NovaSeq platform (Illumina) with mean read lengths of 28bp and 91bp for R1 and R2, respectively. A minimum read depth of 50,000 paired-end reads per cell was targeted for each experiment. Actual read depth ranged from >55,000 paired-end reads per cell to >117,000 paired-end reads per cell, depending on individual experiment and assuming 10,000 cells for each input.

### Single cell analysis

Raw FASTQ files for each experiment were uploaded to the Cloud Analysis platform from 10X Genomics. Read alignment and feature barcode matrices were generated via Cell Ranger Count v6.1.2 (10X Genomics) using standard settings and 10,000 expected number of cells. For all experiments, a custom reference genome was utilized that included unique sequences for tdTomato, ZsGreen, 2A-Cre recombinase, and 2A-Dre recombinase. This reference genome was built following the standard Cell Ranger mkref v3.1.0 pipeline (10X Genomics, https://support.10xgenomics.com/single-cell-gene-expression/software/pipelines/latest/using/tutorial_mr). First, the GRCh38 primary genome assembly along with the comprehensive gene annotation for the primary assembly (v37) were downloaded from the GENCODE website and utilized as inputs to generate the base genome via Cell Ranger mkref. Custom FASTA and GTF files were generated for individual COR-LT system genes, which were then appended onto the base genome as described in the mkref tutorial (10X Genomics).

Complete gene sequences for tdTomato, ZsGreen, 2A-Cre recombinase, and 2A-Dre recombinase that were used to build this custom reference can be found at (https://github.com/lbury3/cor_lt/). In general, sequences include the coding sequence for each gene and the 3’ UTR sequence up but not including the polyA sequence. In one exception, there was a portion of the 3’ UTR sequence that was not included for ZsGreen as it overlapped with the SV40 sequences used as the STOP signal in the reporter constructs. Experimental scRNA-seq data indicated that the upstream CAG promoter drove production of nascent transcripts containing as least a portion of the SV40 polyA sequence in most cells, irrespective of ZsGreen expression (data not shown). Thus, this sequence was excluded from the overall ZsGreen sequence. Removal of this 3’ sequence from the reference genome might lead to under-detection of ZsGreen in scRNA-seq data, as transcripts recovered via the 10X Chromium Next GEM Single Cell 3’ Reagent Kit are enriched in 3’ sequences.

Technical artifacts and background reads were computationally removed via CellBender software,^70^ utilizing the “raw” HDF5 file generated by Cell Ranger as input. Individual parameters were determined by visual analysis of the “Barcode Rank Plot” generated by Cell Ranger and were unique for each experiment (https://github.com/lbury3/cor_lt/). Output files from CellBender where then analyzed in R (v4.1.2) via Seurat software (v4.1.0).^71^ Seurat objects were generated and individual experiments were integrated together following the general pipeline described in the Seurat vignettes (https://satijalab.org/seurat/articles/sctransform_v2_vignette.html). Quality control parameters included upper (6000-7500) and lower (1250) thresholds for the number of genes detected per cell, and an upper threshold for mitochondrial percentage (15% for control and 10-week ASD experiments; 20% for 20-week ASD experiments). Cells outside these parameters were excluded from analysis. Total cell numbers for each experiment after quality control, plus mean nCount_RNA and mean nCount_Features values for each experiment can be found in (Supplemental Table 12, https://github.com/lbury3/cor_lt/). For scRNA-seq experiments utilizing *CTNNB1* isogenic cell lines, doublets were computationally removed via scDblFinder,^72^ which was accessed through the SCP R package (https://github.com/zhanghao-njmu/SCP). Normalization was performed via the SCTransform-v2 function, with mitochondrial percentage regressed out. To avoid clustering driven by expression of COR-LT system components, ZsGreen, TdTomato, Cre, and Dre were removed from the list of genes used as integration anchors. PCA dimensions 1:40 were used for the FindNeighbors and RunUMAP functions, with resolution of 1.5 for FindClusters.

UMAP plots of general cell types (Fig. 1H, Fig. 4A, E) were annotated based on enrichment of canonical gene markers in specific clusters (Supplemental Fig. 3, Supplemental Fig. 6, Supplemental Tables 1, 5, https://github.com/lbury3/cor_lt/). Gene enrichment within individual clusters was determined via the FindAllMarkers function in Seurat (min.pct = 0.25, logfc.threshold = 0.25). If mitochondrial genes, lncRNAs, ribosomal genes, the lncRNA gene MALAT1, and/or lncRNAs in general were enriched in a specific cluster, it was categorized as “Other”, regardless of enrichment of other cell-type-specific genes (Supplemental Tables 1, 5, https://github.com/lbury3/cor_lt/). Non-glutamatergic-neuron-lineage cells (e.g. pericytes, microglia, etc.) were also classified as “Other”. These cells were excluded from downstream analysis.

Gene expression differences between ZsGreen+ and ZsGreen-upper layer neurons in control organoids were determined via the FindMarkers function in Seurat (logfc.threshold = 0, assay = “SCT”). For this comparison, only cells in upper layer clusters where at least one transcript of TdTomato was detected were considered. Cells where at least one transcript of ZsGreen was detected were considered ZsGreen+. Cells where no transcripts of ZsGreen were detected were considered ZsGreen-. ZsGreen+/− comparisons for “cluster 1” were completed in a similar fashion, with only the cells in cluster 1 being analyzed. For randomized comparison of upper layer neurons, two groups of cells were created that were the same size as the previously compared ZsGreen+ (1118 cells) and ZsGreen-(5294 cells). These cells were from the same upper layer clusters used to determine differential gene expression between ZsGreen+ and ZsGreen-upper layer neurons, but assigned to either group randomly, regardless of ZsGreen expression status.

Differentially expressed genes in ASD organoids were calculated via the FindMarkers function in Seurat in a similar fashion as control (logfc.threshold = 0, assay “SCT”, p_val_adj < 0.05). For GSEA analysis, differentially expressed genes between specific groups were generated via FindMarkers function in Seurat, as described above. “Red” neurons were cells where at least one tdTomato mRNA transcript was detected and no ZsGreen transcripts were detected. “Green” neurons were cells where at least one ZsGreen mRNA transcript was detected without any tdTomato transcripts detected. “Yellow” neurons were cells where at least one transcript of both tdTomato and ZsGreen were detected. “No Color” neurons were cells where neither tdTomato nor ZsGreen transcripts were detected. Genes used for GSEA comparisons were ranked based on avg_log2FC values. Actual GSEA analysis was completed via clusterprofiler (v4.6.2).^73^ The compareCluster function was utilized for GO term enrichment visualization across different groups, using default settings.^73^ GO terms were preselected for visualization and p-values were adjusted with the Benjamini-Hochberg correction.

### Data and code availability

All Supplemental Tables and code used to analyze scRNA-seq data, including all CellBender and Seurat analysis, can be found at (https://github.com/lbury3/cor_lt/).

### Tissue fixation and immunohistochemistry

For tissue fixation, organoids were washed 1x in phosphate buffer solution (PBS) before incubation in 4% paraformaldehyde/PBS for 30-45 minutes. Organoids were then washed 3x with PBS and transferred to 30% sucrose/PBS solution overnight. The following day, organoids had sunk to the bottom of the solution and were subsequently embedded in a 50:50 mixture of optimal cutting temperature (OCT) solution (Thermo Fisher) and 30% sucrose before freezing on crushed dry ice. Organoid blocks were either sectioned immediately or stored at −80 degrees until sectioning. 20um tissue sections were generated via cryostat (Leica CM1950) and put on slides (Azer Scientific) which were stored at −20 degrees until IHC labeling.

For tissue labeling, slides were removed from −20°C storage and allowed to come to room temperature. All washes and incubations were performed at room temperature unless otherwise noted. Slides were washed 2x in PBS-T solution (1x PBS + 0.1% Triton-X-100 (Millipore Sigma)) for 5 minutes per wash before incubation in blocking solution (10% donkey serum (Jackson ImmunoResearch) in PBS-T) for 45 minutes. Blocking solution was then removed and replaced with primary antibody solution for overnight incubation at 4 degrees. Primary antibody solution consisted of primary antibodies diluted in blocking solution at antibody-specific concentrations (Supplemental Table 12, https://github.com/lbury3/cor_lt/). After overnight incubation, slides were washed 4x with PBS-T for 10 minutes per wash. Slides were then incubated in secondary antibody solution for 1 hour. Secondary antibody solution consisted of AlexaFluor-conjugated secondary antibodies at 1:500 dilution in blocking solution (Supplemental Table 12, https://github.com/lbury3/cor_lt/). After secondary antibody incubation, slides were washed an additional 3x with PBS-T for 10 minutes per wash. Following these washes, slides were incubated in DAPI solution (Millipore Sigma, 5mg/ml stock, 1:2000 dilution in PBS) for 5 minutes. Slides were then washed 1x for 5 minutes in PBS. Excess liquid was then removed around organoid sections before mounting coverslip (#1.5, Azer Scientific) with Fluoromount aqueous mounting media (Millipore Sigma).

### Microscopy / visualization / image analysis

Live organoid images were acquired on a Leica DMI6000 inverted microscope utilizing either 5x or 10x objectives (5x Leica N Plan, NA = 0.12, 10x Leica PL Fluotar, NA = 0.3). Excitation/emission filters for ZsGreen were BP 480/40nm and BP 527/30nm, respectively. Excitation/emission filters for TdTomato were BP 560/40nm and BP 645/75nm, respectively. Fixed images were acquired via either a Hamamatsu Nanozoomer S60 slide scanner (Fig. 2D, E; Fig. 3B; Fig. 4C, G; Fig. 5C, D), Leica DM2500 epifluorescence microscope (Fig. 1E), Leica DM6000 epifluorescence microscope (Fig. 3B; Supplemental Fig. 4A), or Leica TCS SP8 gated STED 3X confocal system mounted on a Leica DMI6000 inverted microscope (Fig. 1F, G; Supplemental Fig. 1B; Fig. 3C; Supplemental Fig. 7A). All Nanozoomer images were acquired with a 20xobjective (NA = 0.75). Excitation/emission filters for Nanozoomer images were as follows: DAPI ex.=BP 387/11, em.=BP 440/40; ZsGreen ex.=BP 485/20, em.=BP 525/30; TdTomato/AlexaFluor555 ex.=BP 560/25, em.=BP 607/36; AlexaFluor647 ex.=BP 650/13, em.=BP 684/24. All DM2500 images were acquired with a 20x (Leica PL Fluotar, NA = 0.5) objective. Excitation/emission filters for DM2500 images were as follows: DAPI ex.=BP 360/40, em.=LP425; AlexaFluor488 ex.=BP 470/40, em.=525/50; TdTomato/AlexaFluor555 ex.=BP 560/40, em.=630/75. All DM6000 images were acquired with a 20x (Leica PL Fluotar, NA = 0.5) objective. Excitation/emission filters for DM6000 images were as follows: DAPI ex.=BP 360/40, em.=BP 470/40; AlexaFluor488 ex.=BP 480/40, em.=BP 527/30; TdTomato/AlexaFluor555 ex.=BP 546/12, em.=BP 600/40. Images acquired via SP8 confocal microscope utilized either 20x (Leica Plan Apo, NA = 0.75) or 40x (Leica Plan Apo, NA = 1.3) objectives. Fluorophores were stimulated with either a 405nm diode laser for DAPI, or a tunable white light laser for other fluorophores (ZsGreen = 493nm, TdTomato/AlexaFluor555 = 554nm; AlexaFluor594 = 595nm; AlexaFluor647 = 652nm). Tunable emission filters were set as follows: DAPI = 415nm-460nm; ZsGreen = 500nm-525nm; TdTomato/AlexaFluor555 = 565nm-585nm; AlexaFluor594 = 615nm-635nm; AlexaFluor647 = 670nm-715nm. To ensure TdTomato signal does not bleed through when imaging AlexaFluor594 fluorophores, TdTomato+ samples without AlexaFluor594 labeling were imaged with AlexaFluor594 excitation and emission parameters. No bleed-through signal from TdTomato was detected in these images (data not shown).

Microscopy images were processed and merged images were generated via ImageJ software. For quantification in Fig. 2F and Supplementary Fig. 1C, the locations SATB2+ or TBR1+ cells were manually identified in individual IHC images of separate organoids. These locations were then overlaid on the tdTomato or ZsGreen channels of the same images and the expression of individual reporters (positive or negative) at SATB2+ or TBR1+ locations was manually determined. IHC quantification in Figure 4D was performed using a custom-written ImageJ script that determined pixel overlap between IHC channels. Organoid sections from three individual organoids were analyzed per genotype. Statistical comparisons were performed via unpaired student’s t-test for all IHC quantification.

Cell-type UMAP and violin plot figures were generated via Seurat using the DimPlot and VlnPlot functions, respectively. Cluster and single-gene UMAP plots were generated via the dittoSeq R package^74^ with the dittoDimPlot function. Bar plots were generated via Microsoft Excel or MATLAB and volcano plots were generated via the EnhancedVolcano R package.^75^ For volcano plots in Fig. 2G, H, ZsGreen was removed from the dataset, as its significantly enhanced expression in ZsGreen+ cells was an outlier that skewed the x- and y-axes. ZsGreen was also not included in Supplemental Fig. 9A, where it is upregulated in all three groups. All plots were further processed and combined into figures via Adobe Illustrator.

## Supplemental Information

<u>Supplemental Figure 1. Specific and consistent activation of COR-LT system reporters</u>. (A) Live imaging of iPS cell colonies without recombinase transfection (top row), with transient Cre recombinase transfection (middle row), or with Dre recombinase transfection (bottom row). (B) IHC detection of TBR1 (blue) in 9-week-old organoid sections derived from a cell line with pre-activated ZsGreen (green) and tdTomato (red). (C) Quantification of co-localization between TBR1 and either tdTomato or ZsGreen reporters in individual images of pre-activated organoids. Scale bars: (A) = 100um, (B) = 50um.

<u>Supplemental Figure 2. Split UMAPs for individual experiments</u>. (A) Split UMAPs for individual replicates of control scRNA-seq experiments from 4-month-old organoids. (B, C) Split UMAPs for CTNNB1 WT/WT and CTNNB1 WT/Q76* experiments from 10-week-old (B) and 20-week-old (C) organoids.

<u>Supplemental Figure 3. Clustering and cell-type-specific gene expression in scRNA-seq experiments from control organoids</u>. (A) UMAP with cell clusters as determined by the FindClusters function in Seurat, resolution = 1.5. (B) UMAPs indicating expression of cell-type-specific gene markers, including astroglial progenitors (*SOX2*, *HES1*, top row), IPCs (*HES6*, *NHLH1*, top middle row), upper layer neurons (*SATB2*, *EPHA4*, middle row), deep layer neurons (*LMO3*, *PDE1A*, bottom middle row), and cycling progenitors (*MKI67*, *TOP2A*, bottom row).

<u>Supplemental Figure 4. tdTomato expression in neurons of 5-week-old control organoids</u>. (A) IHC detection of tdTomato (magenta) and TBR1 (green) in 5-week-old control COR-LT organoid sections. Scale bars = 50um.

<u>Supplemental Figure 5. Reporter expression and GSEA in four-month-old control COR-LT organoids</u>. (A, B) Violin plots of tdTomato and ZsGreen expression as determined by scRNA-seq in 4-month-old control COR-LT organoids. (C) GSEA between distinct lineages of 4-month-old control upper layer neurons. Top box (“Lineage #1”) indicates GO terms enriched in neurons of the first lineage listed on the x-axis, bottom box (“Lineage #2”) indicates GO terms enriched in neurons of the second lineage listed on the x-axis. For “Yellow vs No Color” and “Red vs No Color” comparisons, GO terms were preselected for visualization and p-values were adjusted with the Benjamini-Hochberg correction. (D) Volcano plot of differential gene expression between upper layer neurons in randomized groups. No genes were differentially expressed at a statistically significant (p_val_adj < 0.05) level. Horizontal dashed line = p_val_adj = 0.05; vertical dashed line = 0.

<u>Supplemental Figure 6. Clustering and cell-type-specific gene expression in scRNA-seq experiments from ASD organoids</u>. (A, B) UMAPs of scRNA-seq experiments from 10-week-old (A) and 20-week-old (B) ASD organoids with cell clusters as determined by the FindClusters function in Seurat, resolution = 1.5. (C, D) UMAPs of scRNA-seq experiments from 10-week-old (C) or 20-week-old (D) organoids, indicating expression of cell-type-specific gene markers, including astroglial progenitors (*SOX2*, *HES1*, top row), IPCs (*HES6*, *NHLH1*, top middle row), upper layer neurons (*SATB2*, *EPHA4*, middle row), deep layer neurons (*LMO3*, *PDE1A*, bottom middle row), and cycling progenitors (*MKI67*, *TOP2A*, bottom row).

<u>Supplemental Figure 7. Verification of Cre recombinase expression in EOMES+ IPCs</u>. (A) IHC detection of EOMES (magenta) and Cre recombinase (green) in 10-week-old CTNNB1 WT/WT (top) and CTNNB1 WT/Q76* (bottom) COR-LT organoid sections. Yellow boxes in first and third rows indicate regions shown in second and fourth rows, respectively. Scale bars: rows 1,3 = 50um, rows 2,4 = 25um.

<u>Supplemental Figure 8. scRNA-seq verification of the COR-LT system in ASD organoids</u>. (A, C) UMAPs of COR-LT component expression in individual cells of 10-week-old (A) and 20-week-old (C) ASD organoids. (B, D) Violin plots of COR-LT component expression within specific cell groups of 10-week-old (B) and 20-week-old (D) ASD organoids.

<u>Supplemental Figure 9. Common genes that characterize neuronal lineage across genotypes</u>. (A, B) Venn diagrams that indicate the number of genes whose expression is enriched in “yellow” lineage upper layer neurons (A) and “red” lineage upper layer neurons (B), when the gene expression of “yellow” and “red” upper layer neurons is directly compared in each group. Control data is from 4-month-old organoids while data from ASD organoids are from 20-week-old organoids. Genes common to all three groups or common to the control and CTNNB1 WT/WT groups are displayed.

