## Supplemental Figures for "Neuronal lineage tracing from progenitors in human cortical organoids reveals novel mechanisms of human neuronal production, diversity, and disease"

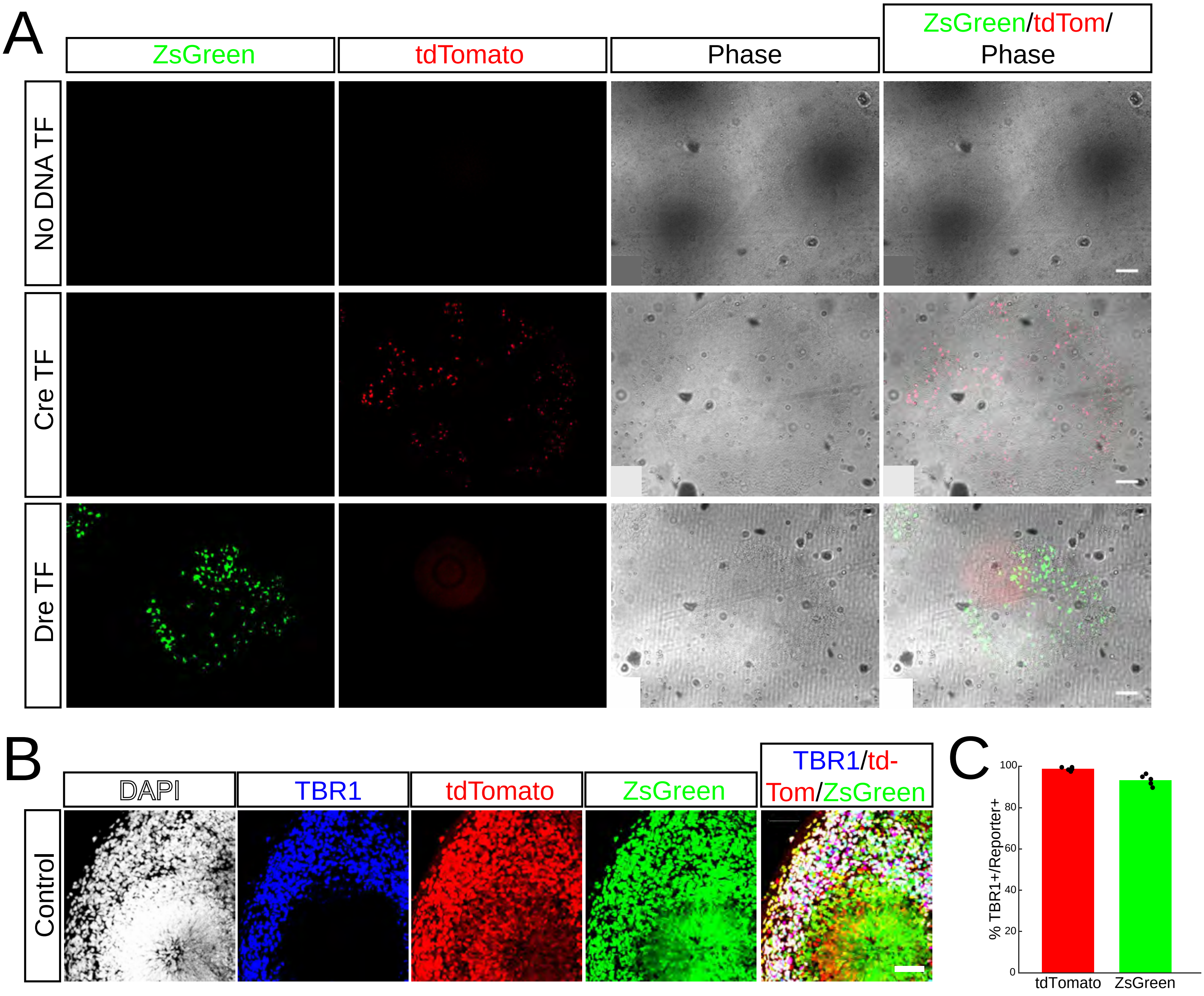


Supplemental Figure 1

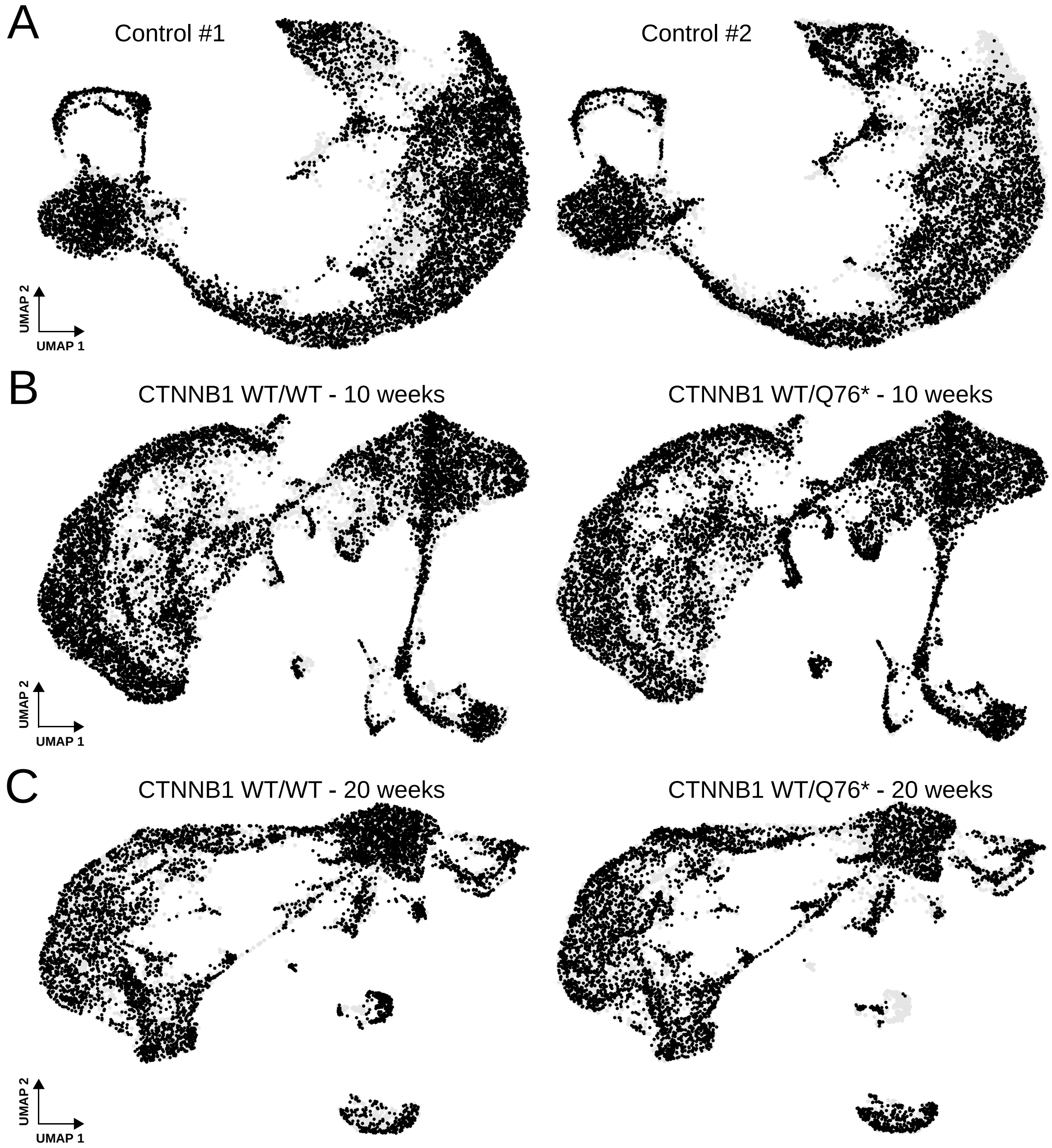


Supplemental Figure 2

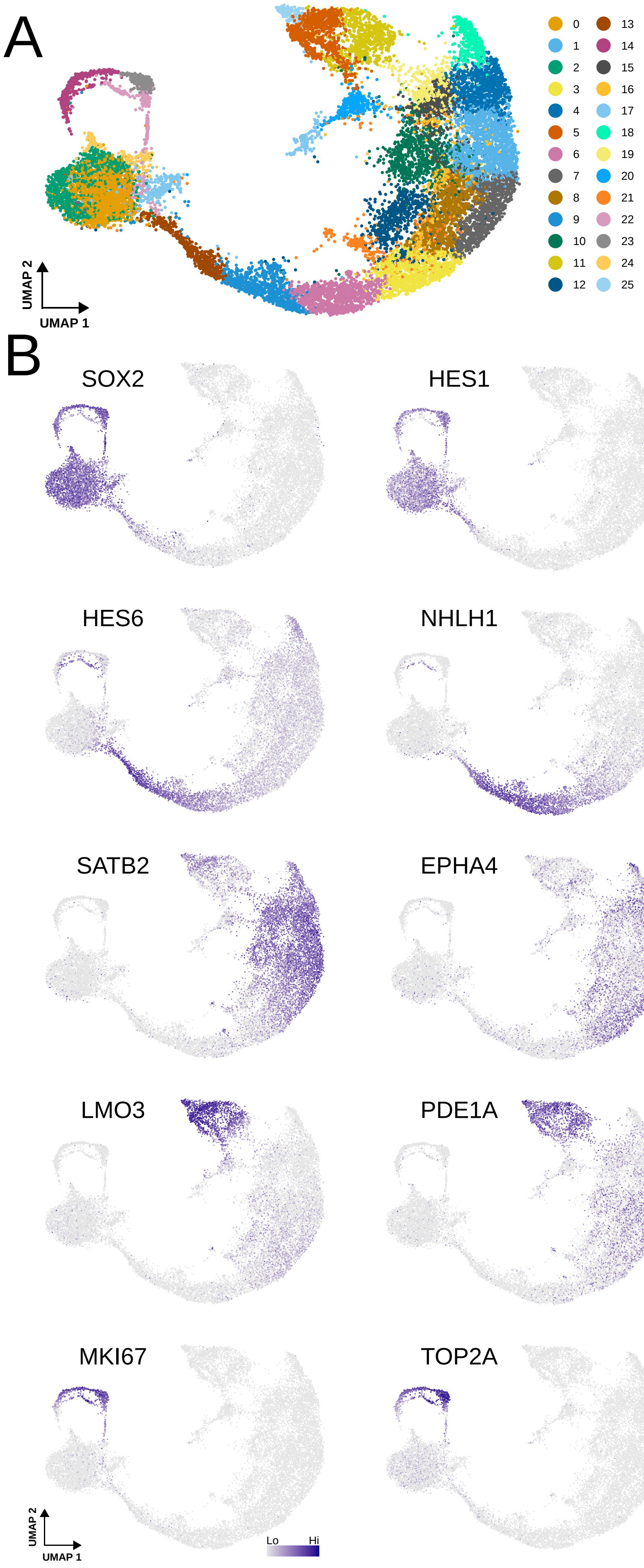


Supplemental Figure 3

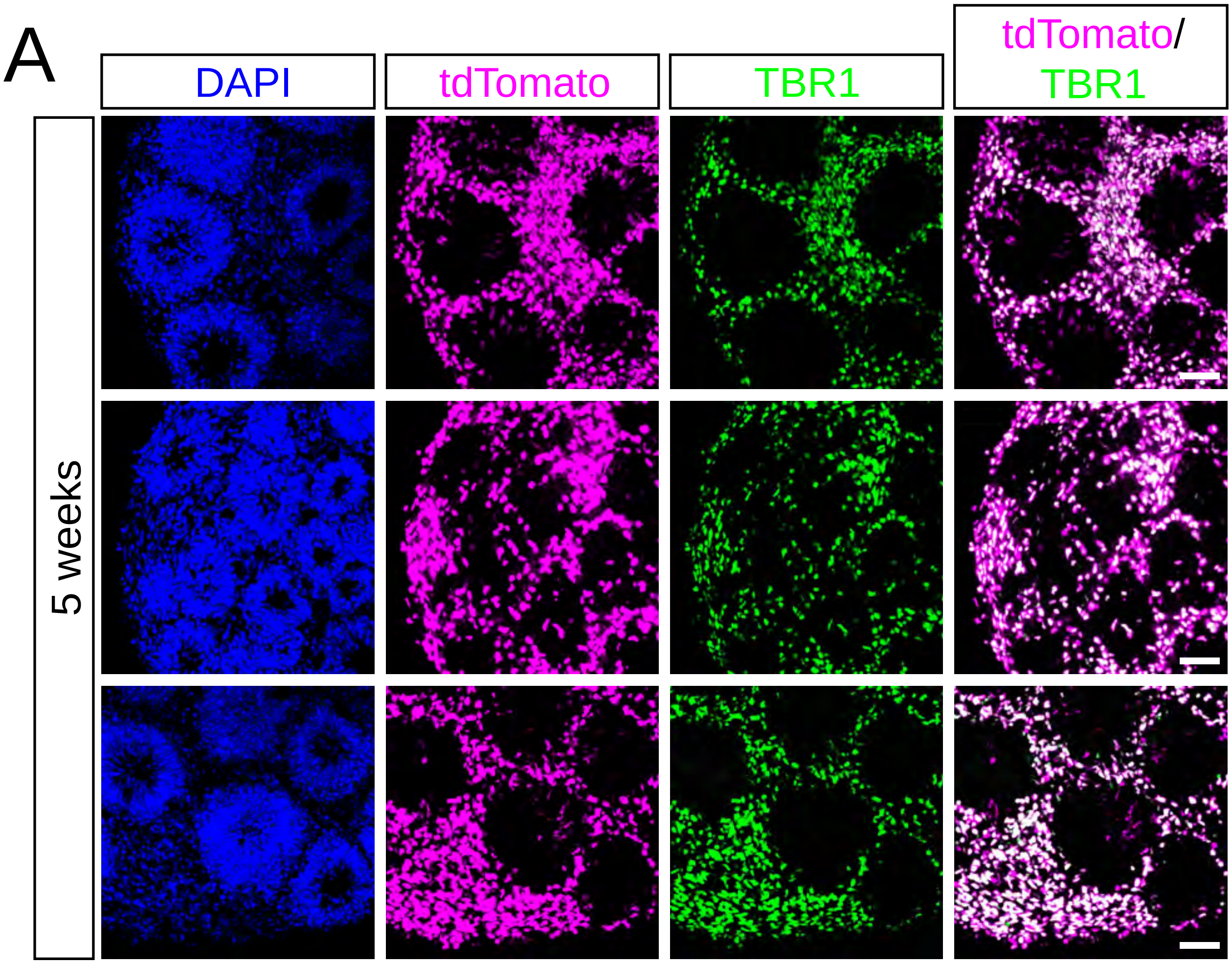


Supplemental Figure 4

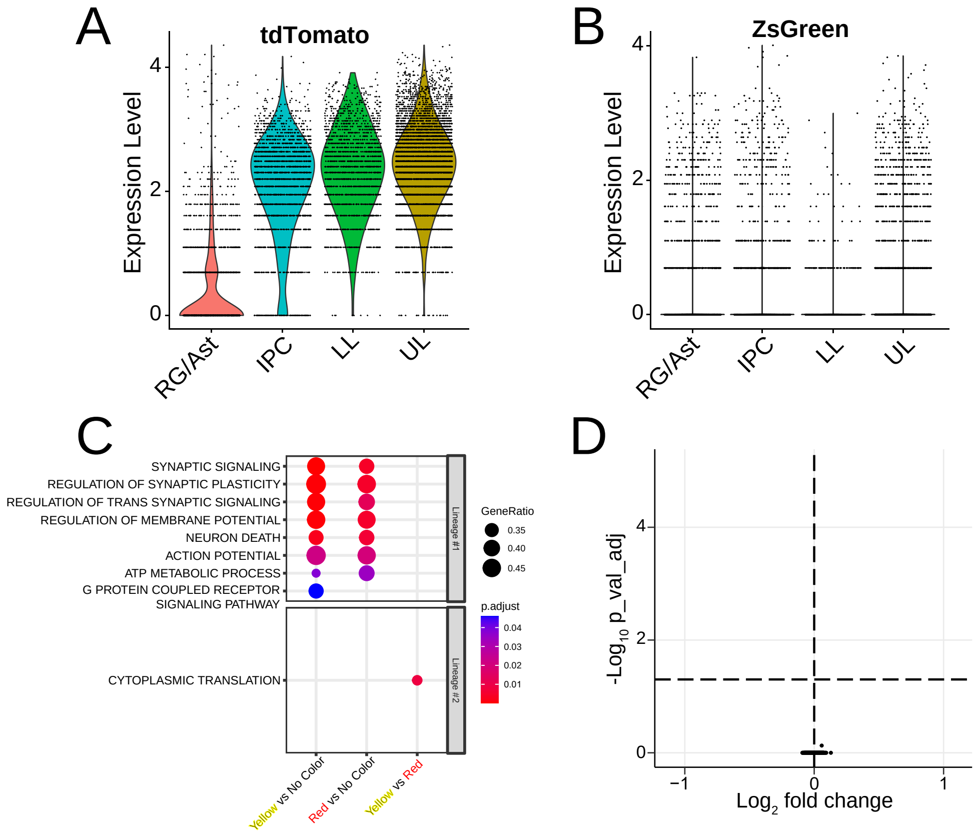


Supplemental Figure 5

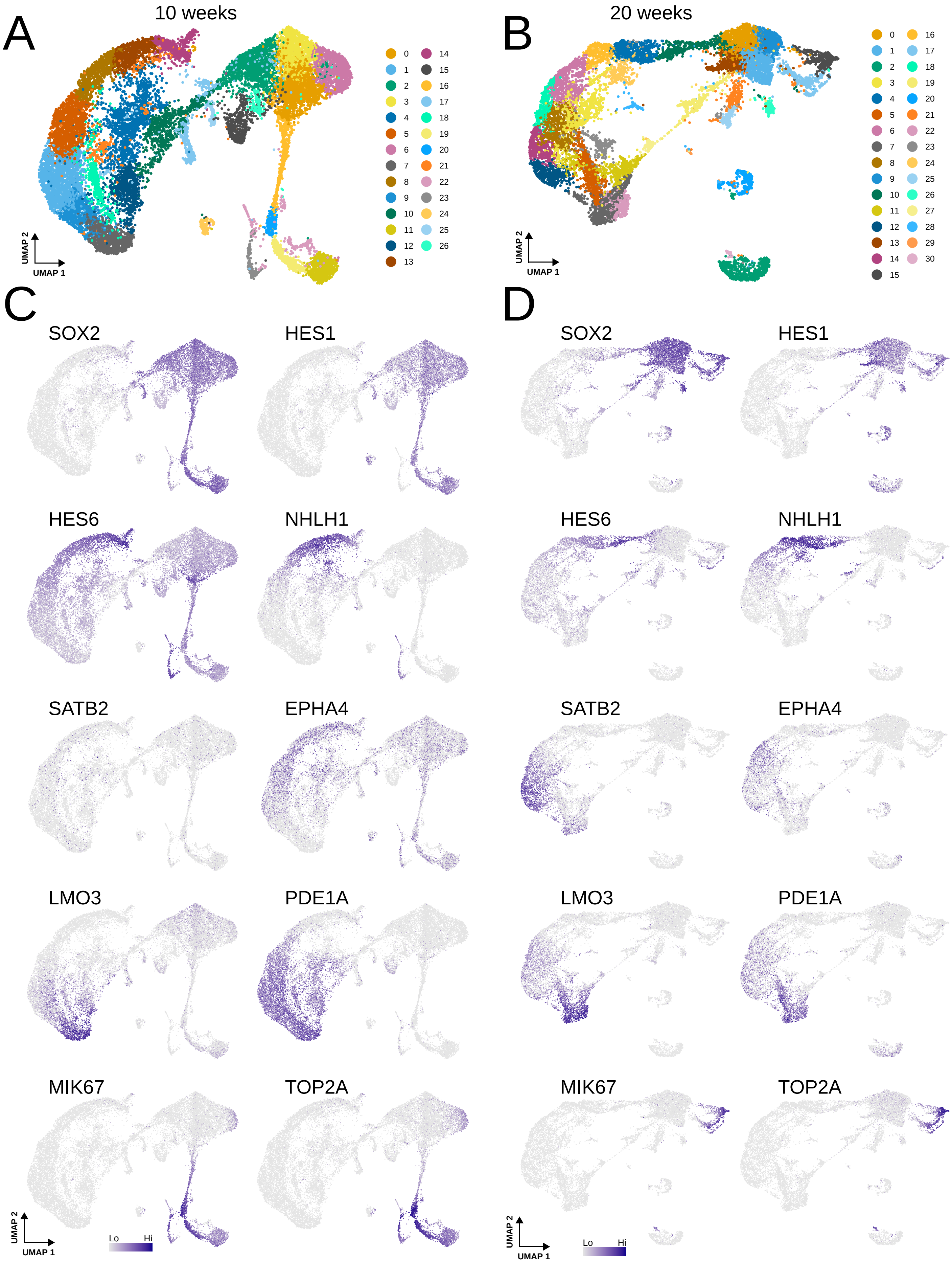


Supplemental Figure 6

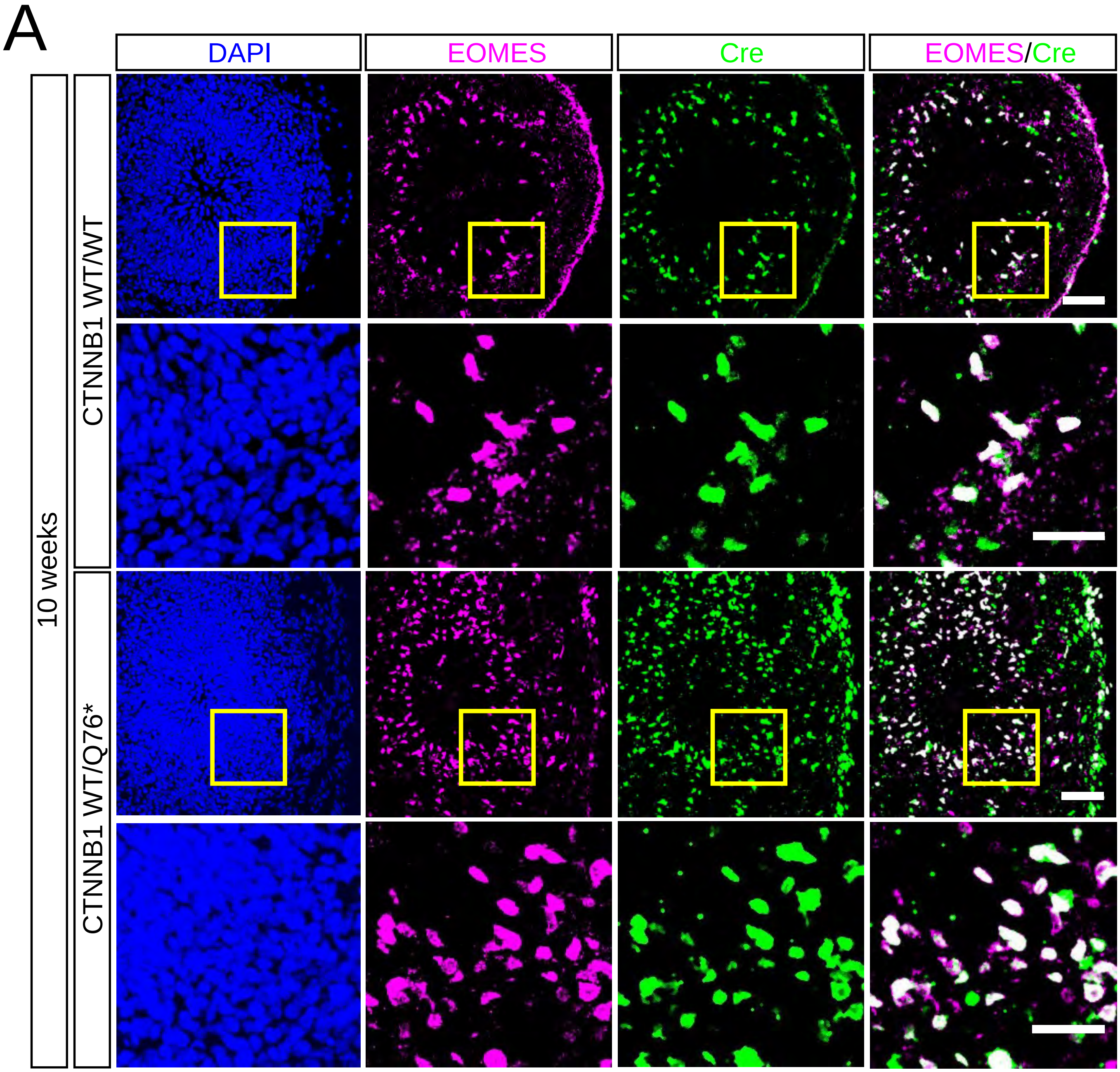


Supplemental Figure 7

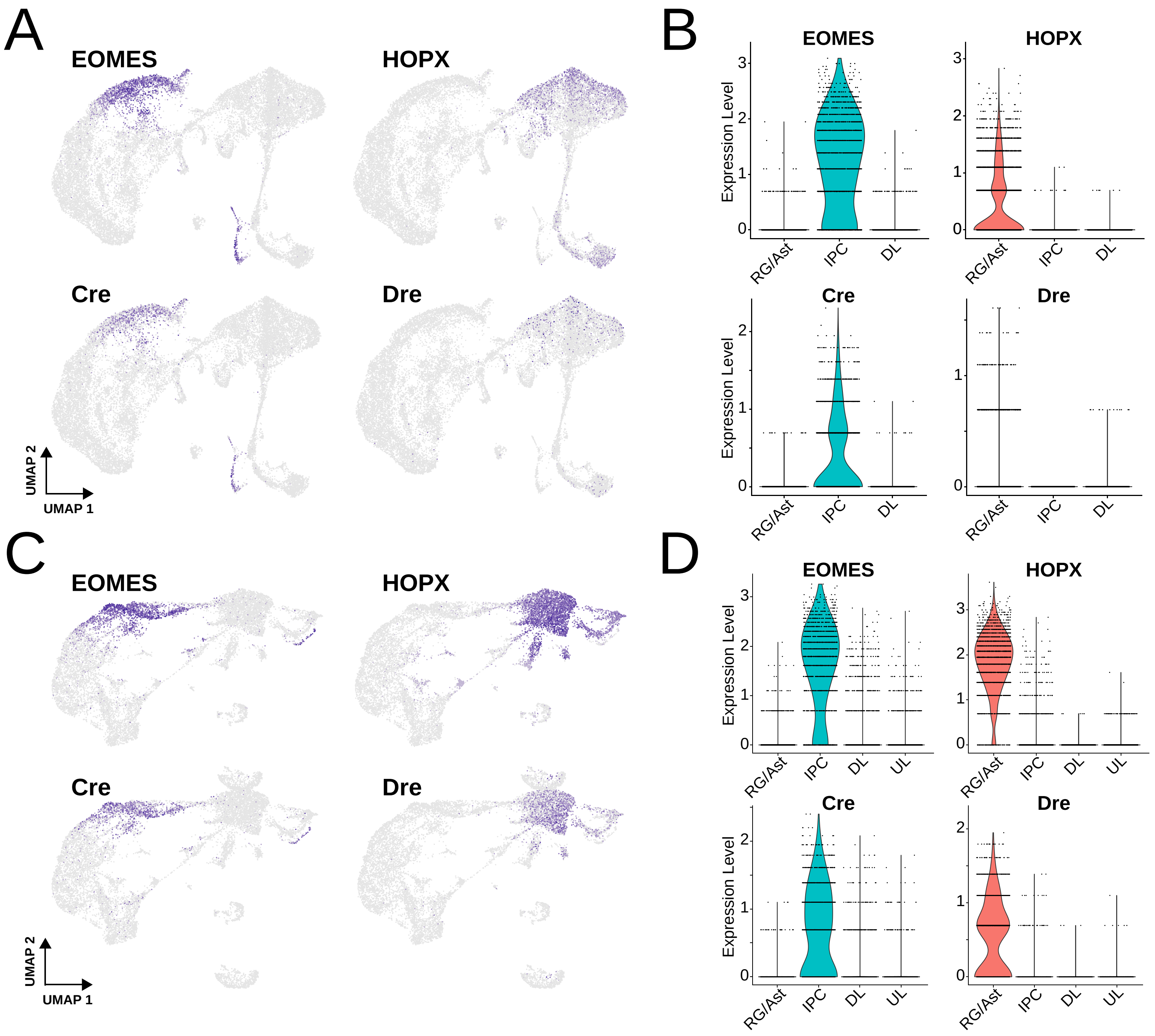


Supplemental Figure 8

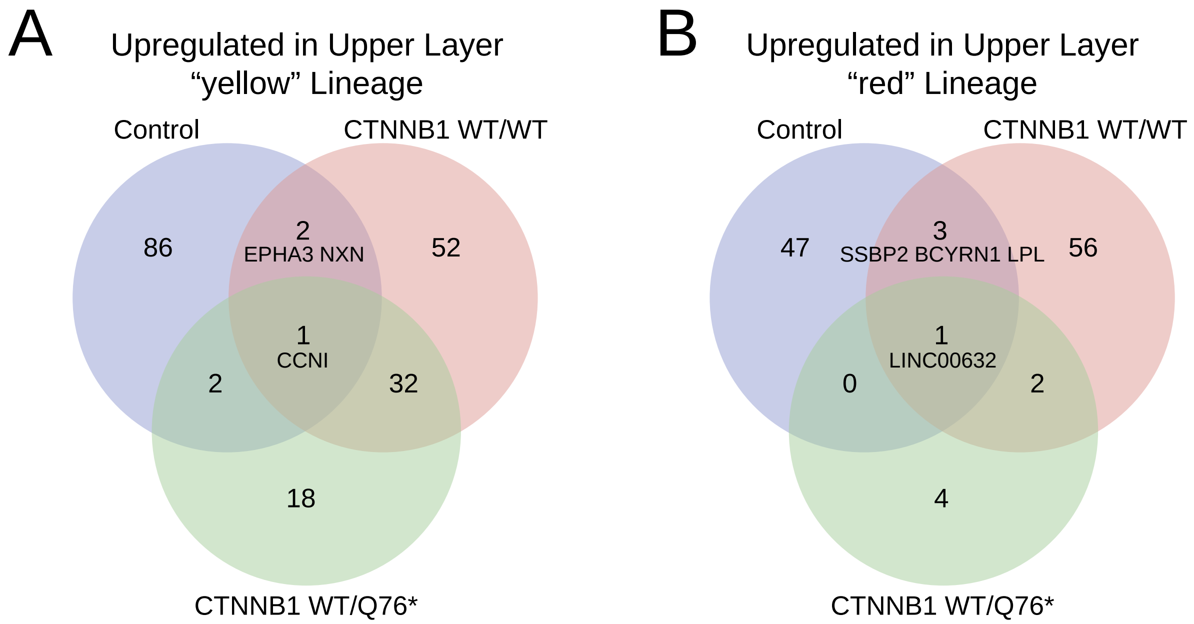


### Supplemental Figure 9
